# Combinatorial mathematical modelling approaches to interrogate rear retraction dynamics in 3D cell migration

**DOI:** 10.1101/2020.08.03.234021

**Authors:** Joseph H.R. Hetmanski, Matt Jones, Fatima Chunara, Jean-Marc Schwartz, Patrick T. Caswell

**Affiliations:** Wellcome Trust Centre for Cell-Matrix Research School of Biological Sciences Faculty of Biology Medicine and Health The University of Manchester Manchester, U.K.

## Abstract

Cell migration in 3D micro-environments is a complex process which depends on the coordinated activity of leading edge protrusive force and rear retraction in a push-pull mechanism. While the potentiation of protrusions has been widely studied, the precise signalling and mechanical events that lead to forward movement of the cell rear are much less well understood, particularly in physiological 3D extra-cellular matrix (ECM). We previously discovered that rear retraction in fast moving cells is a highly dynamic process involving the precise spatiotemporal interplay of mechanosensing by caveolae and signalling through RhoA. To further interrogate the dynamics of rear retraction, we have adopted three distinct mathematical modelling approaches here based on (i) Boolean logic, (ii) deterministic kinetic ordinary differential equations (ODEs) and (iii) stochastic simulations. The aims of this multi-faceted approach are twofold: firstly to derive new biological insight into cell rear dynamics via generation of testable hypotheses and predictions; and secondly to compare and contrast the distinct modelling approaches when used to describe the same, relatively under-studied system. Whilst Boolean logic was not able to fully recapitulate the complexity of rear retraction signalling completely, our ODE model could make plausible population level predictions. Stochastic simulations added a further level of complexity by accurately mimicking previous experimental findings and acting as a single cell simulator. Our approach has also highlighted the unanticipated potential of targeting CDK1 to abrogate cell movement, a prediction we confirmed experimentally. Moreover, we have made a novel prediction regarding the potential existence of a ‘set point’ in local stiffness gradients that promotes polarisation and rapid rear retraction. Overall, our modelling approaches complement each other, suggesting that such a multi-faceted approach is more informative than methods based on a single modelling technique to interrogate biological systems.

## Introduction

Cell migration is a key physiological process which underpins development, immune responses, wound healing, and the metastatic progression of diseases such as cancer. Archetypal mesenchymal motility has been conceptualised as a cyclic process depending on the regulated activity of four key stages (Lodish, 2003; Ridley *et al*., 2003): polarisation to form distinct leading and trailing edges (Lauffenburger and Horwitz, 1996); forward protrusion of the leading edge (Mogilner and Oster, 1996); adhesion to the substratum (Hynes, 1987); and retraction and forward propulsion of the cell rear (Riento and Ridley, 2003). Of these steps, the retraction of the rear remains the most neglected area of study, despite it being essential for fast movement (Cooper and Hausman, 2013; Hetmanski *et al*., 2019). The key instigator of cell rear contractility is the ‘switch-like’ small GTPase RhoA (Ridley and Hall, 1992; Etienne-Manneville and Hall, 2002): RhoA activity, via effector proteins Rho kinase (ROCK1/2) (Riento and Ridley, 2003) and/or PKN-2 (Quilliam *et al*., 1996) leads to phosphorylation of the motor-protein non-muscle myosin II (NMMII), which provides the force required to retract the rear of the cell (Pellegrin and Mellor, 2007); RhoA is further required to activate diaphanous related formins (DRFs) including mDia-1 (O’Connor and Chen, 2013), which are required to reorganise the actin cytoskeleton at the trailing edge in a polymerising capacity (Breitsprecher and Goode, 2013).

Recently, we identified a mechanism for efficient rear retraction in fast moving, highly polarised cells in 3D-matrix and on 2D rigidity gradients (Hetmanski *et al*., 2019). We found that caveolae, mechanosensitive invaginations of the plasma membrane (Parton and del Pozo, 2013), are involved in sensing low membrane tension, which is established at the cell rear in cells polarised by the physical properties of their environment (durotaxis; the preference of cells to move towards areas of stiffer substrate (Lo *et al*., 2000)). Caveolae then propagate these mechanical inputs to locally activate RhoA via the guanine exchange factor (GEF) Ect-2. We further found that RhoA acts primarily through ROCK1 and PKN-2 to organise the acto-myosin machinery into a conformation conducive to highly dynamic rear contractility, while ROCK2 (and other PKNs 1 and 3) are redundant (Hetmanski *et al*., 2019). We produced a detailed cell rear model during 3D migration, incorporating the temporal dynamics of both signalling and mechanical events. Simulations using this model reinforced our hypothesis that efficient rear retraction is a positive feedback process, and the unexpected prediction that perturbation of ROCK1 and PKN-2 disrupts caveolae formation, despite appearing upstream in the model (a prediction subsequently validated experimentally (Hetmanski *et al*., 2019)). Since this was an early rendering of such detailed dynamics and there remain numerous unknowns at play within the network, it is an ideal system to interrogate using three distinct modelling approaches of varying complexity: Boolean logic, deterministic ordinary differential equations (ODEs) and stochastic simulations.

While the dynamics leading to cell rear retraction specifically has not been widely modelled mathematically, other signalling/mechanical events leading to the potentiation of the cell motility machinery have been interrogated successfully using the approaches mentioned above, particularly regarding the formation of leading edge protrusions (Nikonova *et al*., 2013; Kim *et al*., 2015; Byrne *et al*., 2016; Hetmanski *et al*., 2016). Like many cellular processes, efficient cell migration is incredibly complex: the signalling pathways requiring precise, regulated integration in order to polymerise actin at the leading edge appropriately (Hetmanski, Schwartz and Caswell, 2016a; Hetmanski, Schwartz and Caswell, 2016b). Given this complexity, it is unsurprising that mathematical models are increasingly relied upon to organise and formalise knowledge and in turn derive new underlying biological knowledge via the validation/refusal of testable predictions. For example, we and others have previously used simple Boolean logic to investigate large-scale growth factor signalling networks leading to differential Rho GTPase dynamics (Samaga *et al*., 2009; Singh *et al*., 2012; Kim *et al*., 2015; Hetmanski *et al*., 2016); others meanwhile have utilised more quantitative ODE based approaches to investigate the cross-talk of Rac1/RhoA with their immediate upstream regulators in much smaller systems (Tsyganov, Kolch and Kholodenko, 2012; Nikonova *et al*., 2013; Byrne *et al*., 2016).

Generally, particularly in more well-characterised systems with more known *a priori*, the most appropriate modelling approach is employed to fully interrogate the system of interest in each study. For large systems (> 20 variables) with little quantitative time-course data, a Boolean approach is ideal for ascertaining whether the overall model topography is correct and for making predictions regarding the complete removal of variables or alteration of initial conditions (Wang, Saadatpour and Albert, 2012). Due to the constraints that the activity of all variables is binarised to ON or OFF in a Boolean model and that all reactions take the same time increments, the dynamics of certain systems are simply too complex to study with Boolean logic. When the system is small, and/or detailed *a priori* quantitative knowledge is available, kinetic, ODE based models can be of great value (Sible and Tyson, 2007). Their usefulness is exemplified by the more specific predictions possible and the various tool-kits of subsequent analyses available such as parameter sensitivity and steady-state perturbation (Hoops *et al*., 2006). Provided the topology is correct, ODE models can be excellent predictive tools for average population activity. But one major caveat is their lack of inherent randomness and stochasticity: cell migration dynamics vary incredibly both between cells and within the same cell on a temporal basis (Stokes, Lauffenburger and Williams, 1991; Ouaknin and Bar-Yoseph, 2009), behaviour which simply cannot be replicated with deterministic ODEs. To circumvent this, approaches based on rules and probabilistic rates have been developed instead (Tweedy *et al*., 2016; Bangasser *et al*., 2017), for which all simulations can give wildly altering outputs and can thus be conveniently conceptualised as ‘single cell simulators’ to maximise synergy between *in silico* and *in vitro* results.

Using these complementary approaches in tandem, we aim to both elucidate more predictions and underlying biological knowledge regarding the precise spatiotemporal dynamics which lead to efficient rear retraction and identify possible intervention strategies to prevent this. We also aimed to highlight the strengths, potential pitfalls, and points of agreement and of divergence of the varying modelling approaches when studying a mid-level sized network in more general terms.

## Results

### Boolean model outputs depend on reaction formulation and updating scheme

We developed a model of cell rear retraction in 3D environments based on our previous work (Hetmanski *et al*., 2019), augmented by literature mining to include 19 variables of which 12 were individual proteins), 6 were empirical entities pertaining to stiffness, tension, polarisation, actin alignment and retraction itself, the nominal output of interest (Figure 1A), and caveolae simplified to a single node (for detailed model construction, see Methods). The simple wiring diagram of the system is shown in Figure 1A. Before any simulations were performed however, some ambiguities were identified *a priori*, specifically pertaining to variables which are affected by a combination of activators and inhibitors. In particular, F-actin alignment requires active diaphanous related formins (DRF) and active PKN-2, while the capping protein Cofilin opposes F-actin alignment and thus has an inhibitory role. In Boolean logic terms, this can be incorporated into a model in various guises, here termed OR, AND and Hybrid (Figure 1B, see Methods). Similar OR, AND and Hybrid schemes can describe the requirement of active ROCK1 / inactive MLCP for myosin phosphorylation (pMLC activation, Figure 1B). Together with a Hybrid scheme feeding into RhoA (to achieve intermediate RhoA activation, discussed in detail in the next section), this gives 9 combinations of OR, AND and Hybrid schemes for F-actin alignment and pMLC (both of which are required in an AND reaction for rear retraction). Interestingly, simulating these 9 different combinations gave different outputs for rear retraction levels depending on whether the updating scheme used was deterministic synchronous (all possible reactions occur simultaneously at each time point; Figure 1D) or stochastic asynchronous (only one possible reaction chosen randomly based on a uniform distribution occurs per time increment Figure 1C). AND reactions here were simply too stringent to permit rear retraction in deterministic synchronous simulations (Figure 1D, right) as neither ROCK1, PKN-2 and NOT Cofilin nor ROCK1 and NOT MLCP occurred simultaneously. The less stringent OR + OR combination resulted in sustained, steady state retraction (Figure 1D, left), while the combinations involving only Hybrid or Hybrid + OR schemes gave rise to striking cyclic oscillations of retraction activity (Figure 1D, centre). In the unperturbed simulations, at least some retraction should be visible in the outputs (analogous to the rear retraction observed experimentally previously (Hetmanski *et al*., 2019)); therefore if we were to use deterministic synchronous updates only, the conclusions would be that AND schemes are biologically implausible. Remarkably, using stochastic asynchronous updates instead revealed that sustained or cyclic bursts of retraction occurred in the majority of all 9 combinations (Figure 1C), rendering all combinations plausible in this case. These stochastic simulations were in fact more biologically relevant and plausible (Figure 1C, S1): experimentally, cells that are in exactly the same conditions can exhibit vastly contrasting rear retraction efficiency.

**Figure 1:**
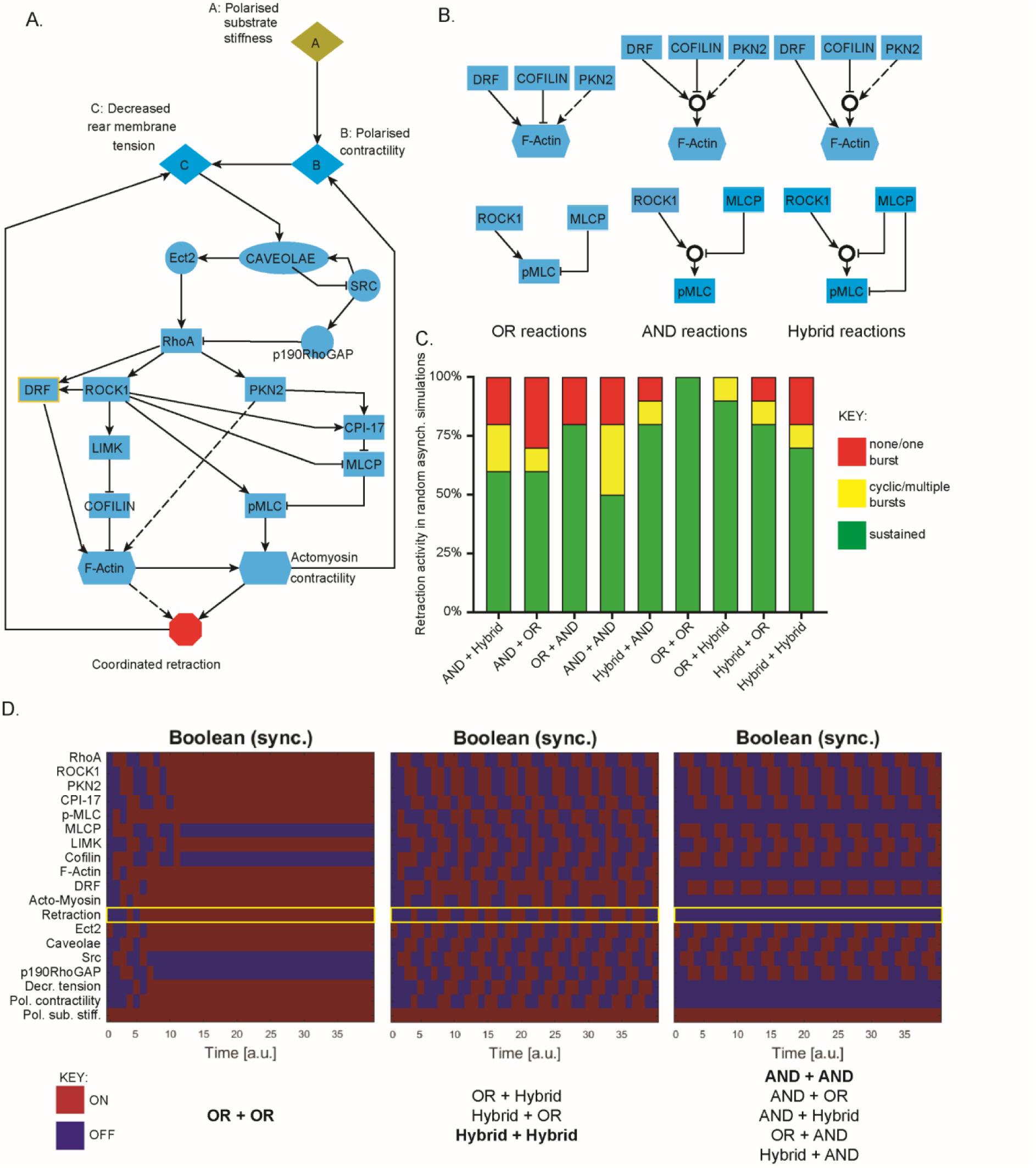
Boolean simulations of cell rear retraction model depend on logical formulation of ambiguous reactions for deterministic synchronous but not stochastic asynchronous updates. A. Model schematic of general variables and reactions used throughout this study. The model comprises of 19 variables of which 12 are individual proteins, 6 are empirical entities pertaining to stiffness, tension, polarisation, actin alignment and retraction itself and the caveolae complex simplified to a single node. B. Schematic representations of different logical inferences used for ambiguous reactions (in which both activator and inhibitor interactions feed into the same node) downstream of RhoA activation: All interactions can independently occur for the OR scheme, reactions require coordinated activity of all inputs for the AND scheme (AND gate depicted as O) and a mix between pure OR and pure AND reactions for the Hybrid scheme as indicated. C. Summary results of the rear retraction activity generated by random asynchronous simulations for the 9 combinations of OR/AND/Hybrid reactions leading to F-Actin alignment and MLC phosphorylation respectively. 10 stochastic simulations were run to 1000 timepoints for each scheme and overall behaviour analysed: either reaching a steady-state with ‘rear retraction’ ON (green), reaching a steady state of ‘rear retraction’ OFF following at most one short-lived burst of retractile activity (red), or at least two bursts of retractile activity throughout the time-course (yellow). D. Heatmaps showing ON/OFF activity of all variables in the model throughout the first 40 timepoints of deterministic synchronous simulations, with the output of interest rear retraction highlighted (yellow box). Simulations either show a transition to steady-state retraction ON (left), oscillatory behaviour of retraction throughout the simulations (centre) or no retraction observable throughout the time-course (right). Each of the 9 reaction schemes as in B demonstrates rear retraction activity which fits into one of these three functional sets, as listed below (bold denotes the precise scheme used for the corresponding heatmaps shown).

### Boolean simulations reveal a paradox of Src/Caveolae dynamics but do not depend on initial conditions

Next, we sought to interrogate the nature of reactions leading to RhoA activity by developing similar AND, OR and Hybrid schemes as in Figure 1B (Figure 2A). Additionally, it has also been reported that upstream of RhoA, caveolae (via Cav1 phosphorylation and oligomerisation (Zimnicka *et al*., 2016)) formation can be increased by Src. Furthermore, caveolae also inhibit Src (via pY14-Cav1 binding to and activating Csk, an inhibitor of Src (Grande-García and del Pozo, 2008)), and Src can in turn activate the RhoA inhibitor p190RhoGAP. We therefore sought to elucidate the importance of this reaction in combination with the AND, OR and Hybrid schemes to give 6 combinations as in Figure 2A. As with reactions downstream of RhoA (Figure 1), deterministic synchronous and stochastic asynchronous updating schemes revealed different output dynamics with regards to the levels of rear retraction (with OR + OR reactions for F-actin alignment and pMLC to enable maximal retraction from this module of the model). While for the deterministic synchronous updates, 5 of the combinations exhibited a steady state of sustained rear retraction (Figure 2C, left, centre), the AND scheme in the absence of the Src → Cav reaction revealed a tendency towards no retraction (Figure 2C, right). Stochastic asynchronous simulations revealed that OR, AND or Hybrid schemes all resulted in dominant retraction when the Src → Cav reaction was included, while all schemes exhibited a biologically implausible lack of retraction without the additional activation of caveolae by Src where more simulations display no retraction than sustained or cyclic bursts (Figure 2B). Based on these stochastic, asynchronous updates, we may therefore conclude that the Src → Cav reaction is crucial for plausible output dynamics.

**Figure 2:**
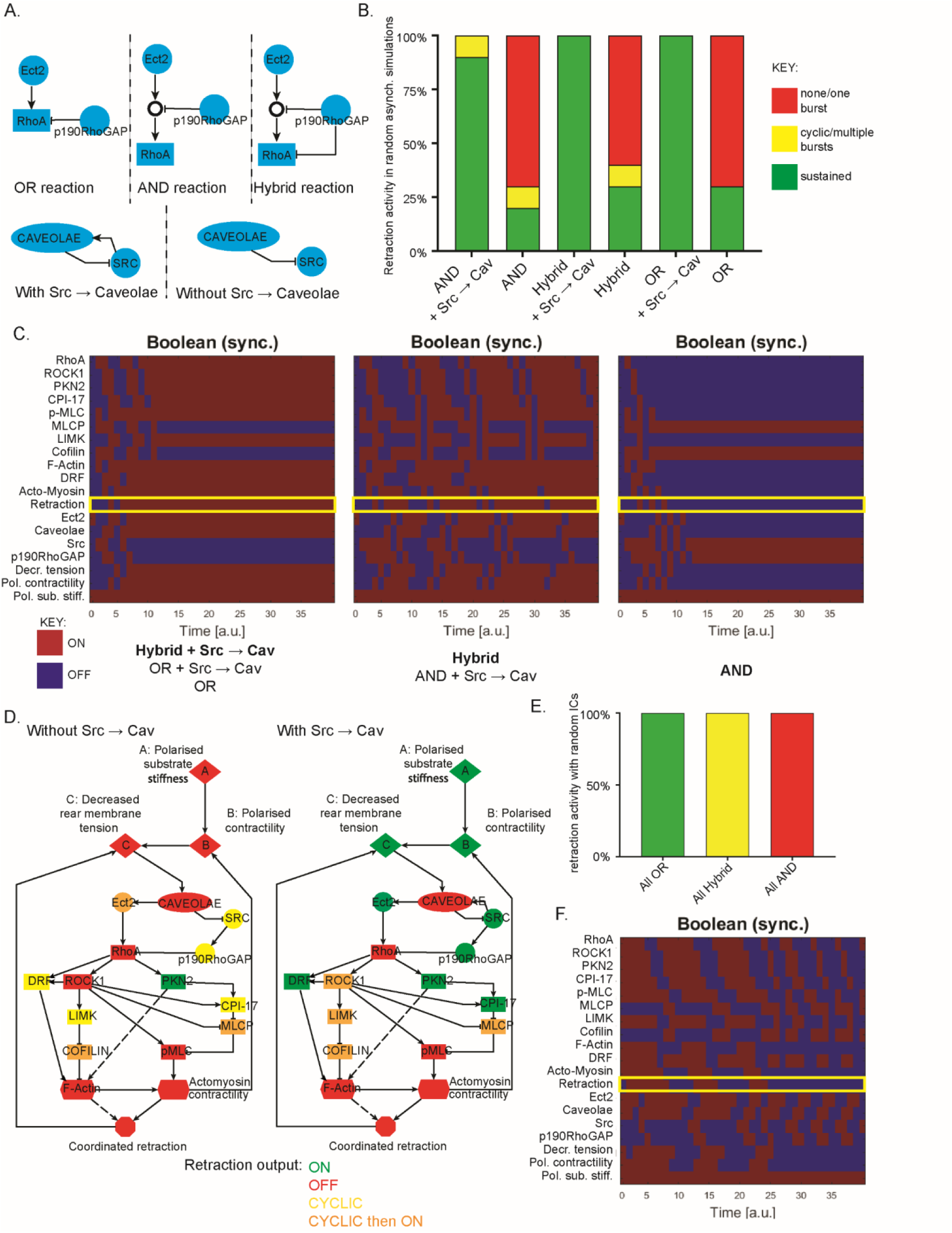
Boolean simulations and *in silico* knockdown predictions depend on Caveolae activation by Src but not on initial conditions. **A.** Schematic summary of OR/AND/Hybrid reactions used upstream of RhoA activation, with and without the ambiguous activation of caveolae by Src. **B.** Summarised outputs of random asynchronous simulations of the 6 different formulations of the model upstream of RhoA activation as in A (with OR + OR reactions used downstream). As in 1C, green denotes steady-state rear retraction activity, red denotes no/one short burst of rear retraction throughout the timecourse and yellow denotes at least two bursts of activity. **C.** Heatmaps showing ON/OFF activity of all variables in the model throughout the first 40 timepoints of deterministic synchronous simulations, with the output of interest rear retraction highlighted (yellow box), with steady state ON retraction reached (left), steady state OFF retraction reached (right) or cyclic bursts of activity throughout the timecourse (centre). The 6 upstream reaction formulations as in A fit into these three functional sets as listed underneath. **D.** Summary of effect on rear retraction output upon individual removal of all variables in the model for the model with (left) and without (right) caveolae activation by Src. Synchronous updates with OR + OR + OR formulation were used (as this gives rise to steady-state retractile activity). *In silico* knockouts could either abrogate rear retraction (red), promote a switch to steady-state oscillatory activity (yellow) or cyclic then ON activity (orange), or result in no change in steady-state rear retraction ON (green). **E.** Summary of rear retraction activity resulting from synchronous simulations with random initial conditions (apart from the input polarised substrate stiffness which is kept ON, and the output retraction which is always initially set to OFF) for the OR + OR + OR, Hybrid + Hybrid + Hybrid and AND + AND + AND schemes as shown. 5 different sets of random ICs were used however steady state behaviour remained unchanged. **F.** Heatmap of the activity of all variables for the synchronous simulation using the AND + AND + AND scheme with the extreme initial conditions of all activator variables ON and all inhibitory variables OFF, showing that a steady-state with rear retraction OFF is still reached.

Next, we sought to make predictions using the Boolean model via *in silico* knockouts. With so many combinations available (6 upstream and 9 downstream giving 54 in total), pure OR + OR + OR schemes with and without Src → Cav subject to synchronous updates were chosen (as these simulations exhibited the desired levels of rear retraction in the unperturbed state and are the least stringent, hence a perturbation effect observed here naturally extends to more stringent combinations and asynchronous updates). Interestingly, perturbing the system in the absence of the Src → Cav directed link revealed predictions which were almost entirely consistent with our previous experimental findings (graphically summarised in Figure 2D, left, S2A): knockout of RhoA, caveolae and ROCK1 or removal of polarised substrate stiffness for example all prevented directional rear retraction *in silico* as strongly suggested *in vitro* previously. Other novel predictions were also rendered which could be experimentally tested in future: for example that LIMK and CPI-17 have a certain importance for rear retraction but are less vital than RhoA/ROCK1. Strikingly, in the presence of the Src → Cav reaction however, many of the predictions seemed at odds with previous experimental findings. Based on our experimental findings, we hypothesized that efficient retraction requires caveolae to sense local decreases in membrane tension and to propagate this to RhoA activity in a positive feedback mechanism. With the Boolean formulation of Src → Cav (alongside Cav ⊣ Src) however, essentially this positive feedback loop is bypassed to hyper-activate and keep caveolae ON, which in turn keeps RhoA ON. This meant that caveolae became the only vital variable upstream of RhoA, while RhoA itself was now the only other protein which when removed abrogates F-actin alignment or pMLC activity and subsequent retraction (Figure 5D, right). This is biologically implausible and implies that the Src → Cav should not be included in the model hence giving rise to a Src*/*Cav paradox: asynchronous updates suggest the presence of the reaction is vital while *in silico* knockouts (with synchronous updates) suggest that it is vital the reaction is not included.

**Figure 5:**
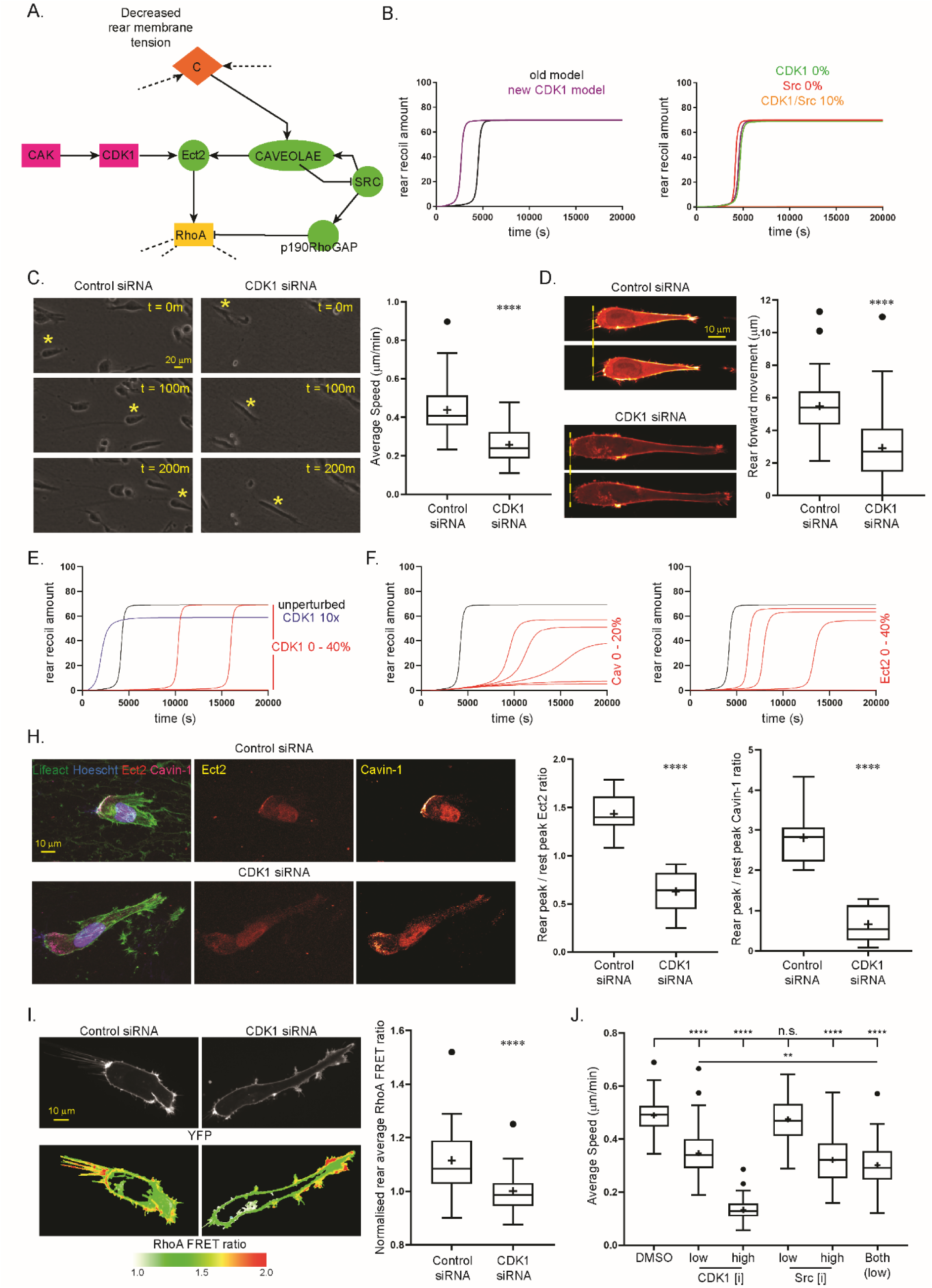
Activation of Ect-2 by CDK1 improves model simulations of rear retraction. **A.** Alternative formulation of model upstream of RhoA, where Ect-2 may be activated by CAK/CDK1 signalling as reported in literature. **B.** Left: Comparison of time course simulations of the retraction amount during the first 20,000 timepoints in the new ‘CDK1’ model (purple line) compared to previous model as in Figure 3 (black line); Right: Time course plot of rear retraction amount in response to complete removal of CDK1 (green line), Src (red line) or knockdown of both to 10% levels (orange line near x-axis) in comparison to the unperturbed simulation (purple line). **C.** Left: A2780 cells were transfected with control (left) or CDK1 (right) siRNA and seeded in CDM and imaged by high-end widefield microscopy across 16 hours, representative individual cells highlighted by yellow * across t = 200 minutes, Right: Quantification of average speed of control or CDK1 knockdown cells during 16h time course, (N = 90 cells across 3 repeats analysed per condition). **D.** Left: Control (left) and CDK1 knockdown (right) A2780 cells expressing Lifeact-Emerald were seeded in 3D CDM for 4 hours and imaged by spinning-disc microscopy for 5 minutes time courses to assess forward rear membrane translocation (yellow dotted line indicates rear position at t = 0); Right: Quantification of average forward rear movement during 5 minute time course of control and CDK1 siRNA cells (N > 50 cells across 3 repeats analysed per condition). **E.** Time course plots of rear retraction amounts in response to *in silico* perturbation of CDK1 with caveolae activation by Src rate appropriately reduced, where red curves show CDK1 activity between 0 40%, blue curve shows 10x overexpression (black curve displays the unperturbed simulation, note values 0, 10 and 20% show negligible retraction and run near the x-axis). **F.** Time course plots of rear retraction amounts in response to *in silico* knockdowns of caveolae between 0 and 20% (left) and Ect-2 between 0 and 40% (right). **H.** Left: Control (top) and CDK1 knockdown (bottom) cells were seeded in 3D-CDM for 4 hours, fixed with PFA and stained for endogenous nuclei (Hoescht), F-Actin (phalloidin), Ect-2 and Cavin-1, maximum intensity projections of whole cell shown for the merged, cavin-1 and Ect-2 channels; Centre, right: quantification of rear peak / leading edge peak Ect-2 intensity (centre) and Cavin1 (right) levels of control and CDK1 knockdown cells (N>10 cells per condition across 2 repeats). **I.** Left: Control (left) and CDK1 knockdown (right) A2780 cells expressing Raichu-RhoA were seeded in CDM and imaged by spinning-disc microscopy to reveal RhoA activity via ratiometric CFP-YFP / YFP-YFP (bottom, LUT shown below, red pixels denote high activity); Right: Average ratiometric RhoA activity at the cell rear of control and CDK1 siRNA cells across 5 minute time lapses, normalised to average CDK1 activity per experimental repeat, N = 31 cells analysed across 3 repeats. **J.** A2780 cells were seeded in CDM and treated with: DMSO (vehicle control); CDK1 inhibitor at low (1 µM) or high (10 µM) dose; Src inhibitor at low (0.3 µM) or high (3 µM) dose; or both CDK1 and Src inhibitors at low (1/0.3 µM) together; 30 minutes prior to capture and imaged by high-end widefield microscopy across 16 hours; Average speed of cells in each condition shown (N = 70 cells across 3 repeats analysed per condition). n.s. denotes non-significant, ** denotes p = 0.01, **** denotes p < 0.0001 by unpaired student t-test or one-way ANOVA.

We further sought to interrogate the effect of the initial conditions (ICs) used on synchronous outputs: previously (Hetmanski *et al*., 2016) we modelled a system based on growth factor initiation, therefore the ICs were obvious. With the rear retraction system however, beyond the controllable input of polarised substrate stiffness, the ICs which lead to efficient retraction are unknown (other than at least one variable in the cascade must be initially active to potentiate the positive feedback mechanisms). In all simulations up to this point, the ICs have been polarised substrate stiffness and Ect-2 ON (see methods). To ascertain whether this was having a stark effect on the synchronous simulation outputs, we randomly assigned ON/OFF activity to all variables in the model (apart from the input node Polarised substrate stiffness (Pol. Sub. Stiff) constrained to ON, and the output node retraction constrained to OFF) before testing the outputs in the AND + AND + AND, OR + OR + OR, and Hybrid + Hybrid + Hybrid combinations. Reassuringly, all sets of random ICs gave rise to exactly the same steady-state dynamics: no retraction with the AND schemes, cyclic activity with the Hybrid schemes and sustained retraction with the OR schemes (Figure 2E, S2B). Moreover, we tested the most extreme case of ICs in which every variable (apart from the output retraction and inhibitory modules) was initially set to ON: again, following some short term bursts of activity, the AND scheme lead to a no retraction steady state (Figure 2F). This data suggests that steady-states are invariant to ICs and no artefacts in the synchronous updates are caused by the ICs used.

Taken together these data suggest the following conclusions can be drawn from the various simulations of the Boolean model:

1. Dynamics of key variables RhoA, F-actin alignment and p-MLC depend partially on the precise balance between their immediate upstream activators and inhibitors in a manner which cannot be fully described using Boolean logic alone.
2. This formulation of the model relies on some level of Caveolae activation by Src, however Boolean logic cannot incorporate this interaction in a way which agrees with previous experimental findings with regards to both unperturbed retraction amounts and knockdowns.
3. Initial conditions are not important to the steady state reached by the model.
4. Incorporating stochasticity into the simulations via random asynchronous updates alters outputs and in many cases leads to more agreement with previous *in vitro* results, suggesting stochastic kinetic models may be of greater value.

As highlighted in points 1 and 2, while Boolean logic can model some aspects of the signalling/mechanical events which lead to efficient cell rear retraction, more quantitative approaches to incorporate the balance of certain reactions are required to precisely model the signalling network that controls rear retraction.

### ODE formulation of rear retraction encapsulates key, known features

Having established that, while the general topography of the Boolean model may be correct, simple ON/OFF logic is unable to fully describe all features of the signalling/mechanical events which lead to efficient cell rear retraction *in vitro*, we increased *in silico* complexity by adopting a kinetic approach based on ODEs with mass action reactions. Generation and details of the modelling approach adopted are described in the methods section and were initially informed by previous findings of the Boolean model. e.g. that the caveolae activation by Src reaction, identified as a potential ambiguity previously, should be included at an appropriately scaled rate (orders of magnitude slower) in comparison with caveolae activation by decreased membrane tension.

We previously used a similar kinetic ODE model to reveal a positive feedback loop centred on caveolae sensing decreased membrane tension and activating RhoA to potentiate real retraction (Hetmanski *et al*., 2019). In the updated model, we included an extra feedback reaction from actomyosin contractility to polarised contractility because we considered that an increase in local contractility would feed into the network in the same way that the rigidity gradient is translated into differential membrane tension. Keeping the rest of the model unchanged, we sought to make further use of deterministic simulations to interrogate cell rear dynamics. The unperturbed time-course simulation exhibited a phase of negligible rear retraction with a very slow increase in speed for the first ∼ 4000 seconds, then a phase of rapid acceleration whereby it takes less than ∼ 1000 seconds for retraction to increase to maximal levels, before a 3^rd^ and final phase of steady state high rear retraction (see black lines in Figure 3A, see S3 for unperturbed time course simulations of all other active variables in the model). This fits previous experimental observations: cells spread, polarise and organise their molecular machinery appropriately (e.g caveolae formation) before moving rapidly in the desired direction and continuing to do until affected by external alterations (such as collision with a neighbour).

**Figure 3:**
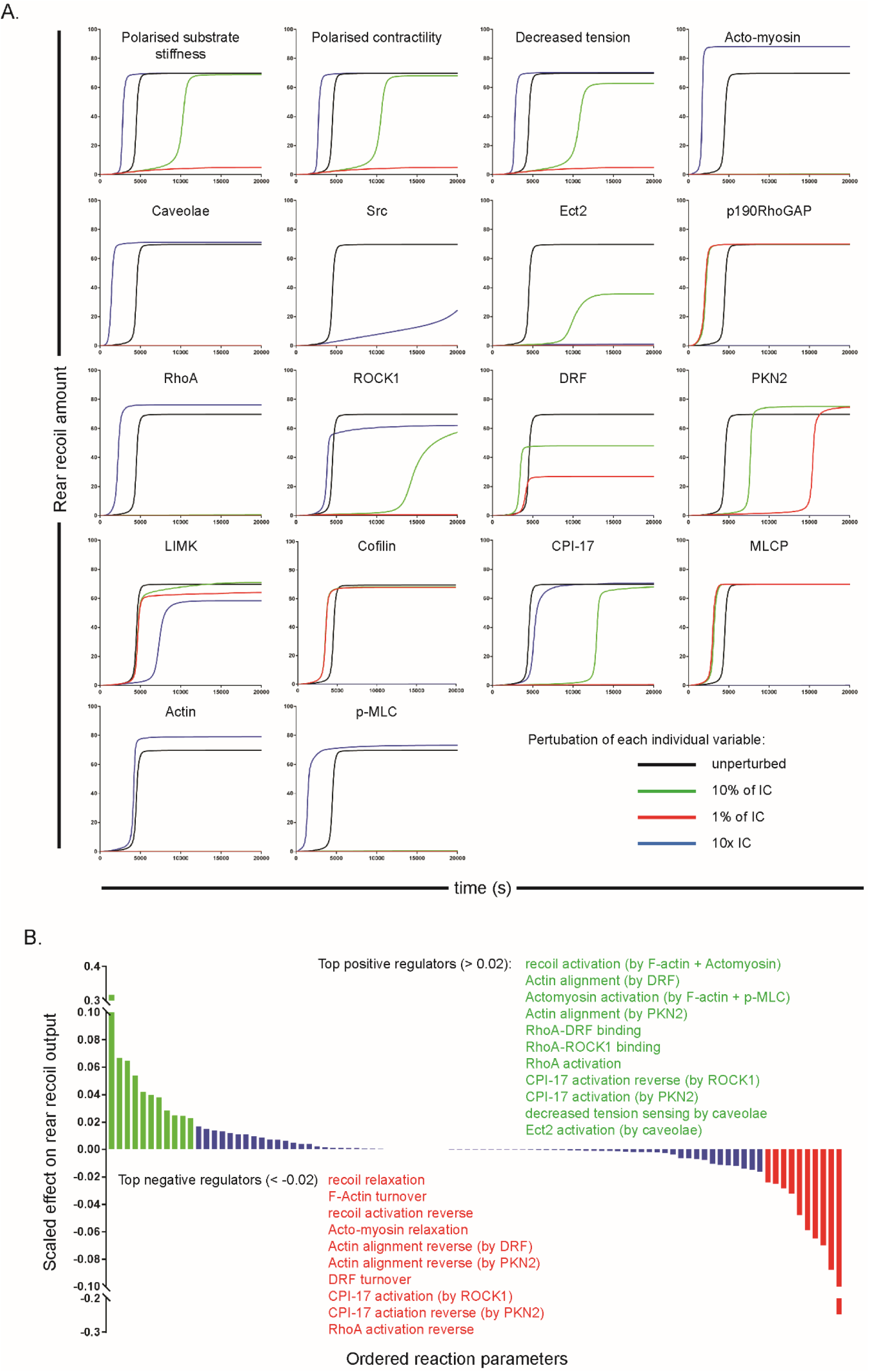
Simulations of the kinetic ODE model reveal variables and reactions which are vital for efficient rear retraction via *in silico* knockdowns and overexpression. A. Time course graphs of the effect on ‘rear retraction amount’ following perturbation of all variables in the model: Each variable was knocked down to 1% level (red line), 10% level (green line) or overexpressed 10x (blue line), while the unperturbed simulation is also shown in each graph for reference (black line). Simulations were run to 20,000s, note lines which appear missing are too close to the origin to be visible (i.e. showing negligible levels of rear retraction). **B.** Summary graph of the parameter sensitivity analysis showing normalised effect of all rates in the model on the rear retraction output. Parameters which have > 0.2 positive effect on rear retraction are shown in green, parameters which have < 0.2 negative effect shown in red, while all other parameters with less drastic effects are represented by the blue bars.

We sought to interrogate the model by systematic perturbation of the amount of every variable in the model by simulating knockdowns to 10% (green lines) and 1% (red lines) levels as well as 10x overexpression (blue lines) compared to unperturbed simulations (black lines) (Figure 3A). This library of individual perturbations agreed with previous experimental observations while also highlighting some potentially interesting features of the model and, plausibly, of the biological processes leading to efficient rear retraction. The inhibitory variables all showed potent abrogation of all rear retraction when 10x overexpressed, while knockdown of the inhibitory proteins results in faster transition to the steady state of high rear retraction speed. Knockdowns of all other variables at 1% or both 1% and 10% had some negative effect on rear retraction to varying levels, affecting either the transition time to steady state or the steady state level. In particular, even at the 10% level, knockdown of RhoA, caveolae, actin and p-MLC all reduced rear retraction to negligible levels, highlighting these proteins (or protein complexes) as ‘bottlenecks’ in this model and thus fundamental regulators of cell rear retraction. This bottleneck observation is further evidenced by the overexpression of all these proteins prompting a faster transition to steady state and an increase in retraction speed at this steady state. The controllable input ‘polarised substrate stiffness’ showed a switch-like response to perturbation: knockdown to 1% (conceptualised as, for example, a near uniform substrate) reduced directional rear retraction to levels 10% of the unperturbed case – which is plausible since cells do indeed retract on uniform substrates, just in a directionally random manner. Knockdown to 10% of polarised substrate stiffness slowed the transition to high rear retraction, but barely reduced this steady-state value.

the 1%, 10% and 10x overexpression simulations of ROCK1, PKN-2 and DRF reveal that efficient rear retraction relies on the precise balance of RhoA binding to effectors. *In silico* DRF knockdowns showed a graded response of rear retraction speed; ROCK1 knockdowns showed more switch-like responses of rear retraction, whereby knockdown to 1% levels abrogates all rear retraction while 10% levels exhibit a severe retardation of transition to high steady-state, but a low decrease in this steady-state value. Knockdown of PKN-2 resulted in a graded increase in transition time to steady state (at the 1% level a marked ∼4 fold increase) but also a slight increase in steady-state rear retraction speed (since the transition time effect is more dominant however, it could still be argued that PKN-2 overall decreases *average* recoil speed, as previously observed). Contrary to the logical expectation that overexpression will have the opposite effect to knockdowns, *in silico* overexpression to 10x levels of DRF, PKN-2 and, to a much lesser extent, ROCK1 all reduced rear retraction. This can be explained by the fact that RhoA acts through these 3 pathways to promote activity of 2 outputs: F-actin alignment and MLC phosphorylation. Overexpression of any one of these effectors however sequesters RhoA away from the other 2 pathways, which leads to a deficiency of either F-actin alignment or p-MLC, thus preventing rear retraction. Finally, as with the Boolean model predictions, Src activating caveolae formation was deemed an important interaction since either knockdown level of Src renders rear retraction speed negligible; overexpression of Src meanwhile also significantly reduced rear retraction speed via the inhibitory effect on p190RhoGAP.

A further method to interrogate the dynamics of the cell rear model was with sensitivity analysis to determine the key kinetic parameters which had the strongest scaled effect on the rear retraction output (Figure 3B). Upon this parameter scan, the first observation was that every parameter in the model has at least some non-zero effect on the rear retraction, highlighting that there is no redundancy in the model formulation. While many reactions had little effect (blue bars, Figure 3B), 11 parameters had a large positive effect on rear retraction speed (> 0.2, the nominal cut-off value and significantly higher than more mundane reactions, green bars) while 10 parameters had an analogous negative effect on our nominated output of interest (red bars). These 21 key parameters exhibited good coverage of the model and further supported the key features required for efficient rear translocation as found previously. The least expected findings of the parameter scan concerned the importance of CPI-17 activation: activation by PKN-2 was important to increasing retraction speed, while surprisingly activation by ROCK1 has a negative effect on rear retraction. This was because ROCK1 is the only activator of p-MLC and LIMK as well as being an activator of DRF, and thus again CPI-17 binding to ROCK1 sequesters the key RhoA effector away from other important functions (an observation which again crucially relies on a finite amount of ROCK1 in the system).

### Further interrogation of key features arising from ODE model predictions

The systematic perturbations of the ODE model in Figure 3 and the Boolean predictions in Figures 1 and 2 highlighted some interesting features of the cell rear retraction model which promoted further examination: in particular the graded versus switch-like perturbation predictions of different variables in the model; the balance between caveolae activation by decreased membrane tension with direct caveolae activation by Src; and the equilibrium between RhoA binding partners. As previously seen (Figure 3A), reducing the polarisation amount of the substrate stiffness to 1% of the original value severely inhibited rear retraction, whereas reducing to 10% merely lengthened the transition time to almost the same steady-state speed of retraction. Further simulations with varying substrate stiffness gradients revealed that there exists a critical switch point of ‘polarised substrate stiffness’ at ∼ 3.9%, below which cells were theoretically unable to directionally retract the rear efficiently, and above which eventually ‘cells’ exhibited rear retraction at the unperturbed steady state speed (Figure 4A). Other variables in the model exhibit similar switch-like dynamics, including proteins such as ROCK1. Other variables, particularly proteins, meanwhile showed a graded response in rear retraction depending on knockdown amount, notably RhoA, caveolae, Ect-2, and DRF. Between the 0% and 50% level of DRF, there was a relatively linear relationship between DRF amount and rear retraction speed (Figure 4B). Again, this rendered the testable prediction that using a DRF inhibitor (such as SMIFH2 for example (Blanchoin and Boujemaa-Paterski, 2009)) at different (low) levels will abrogate retraction to result in a range of cell speeds.

**Figure 4:**
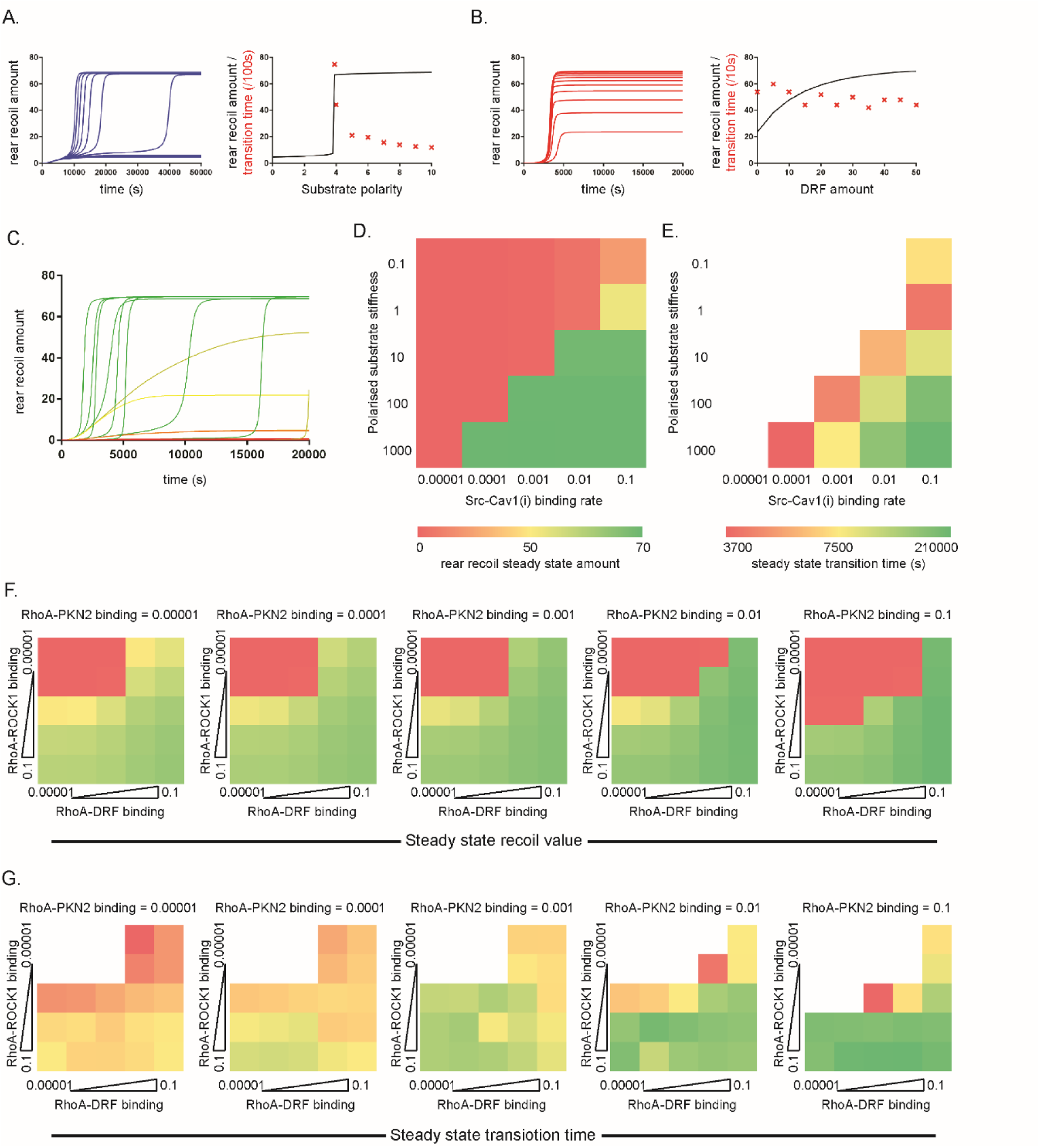
ODE simulations predict switch-like or graded responses to perturbation of different variables and are dependent on the balance of Caveolae activation and RhoA binding to effectors. **A, B.** Time course dynamics of rear retraction amount during the first 50,000s given different **A.** substrate stiffness polarisation amounts (relative to the unperturbed 100% case), with curves corresponding to values ranging from 0 to 10% and **B.** DRF amounts ranging from 0% to 50% (left); steady state rear retraction amount (black line) and transition time to steady state (red x) dependent on **A.** polarised substrate stiffness or **B.** DRF levels (right). **C.** time course plots of the rear retraction amount for the first 20,000s, where each curve represents a different combination of substrate stiffness polarisation and Src-Caveolae (inactive) binding rate. Colours of curves correspond to heatmap summary in D. **D, E.** 5×5 discrete logarithmic heatmaps, displaying how **D.** the steady state rear retraction level and **E.** the transition time to this steady state (if relevant) depend on the combinations of substrate stiffness polarity (Y-axis) and Src-Caveolae binding rate (X-axis). Green denotes high rear retraction / fast transition time, red denotes low / negligible rear retraction / slow transition time as indicated, white squares in E correspond to squares with low steady state rear retraction, thus transition time is not relevant. **F, G.** 5×5 discrete logarithmic heatmaps showing the dependence of **F.** the steady state rear retraction level and **G.** the transition time to this steady state (if relevant) on the binding rates of RhoA to its three effectors: ROCK1 (Y-axes), DRF (X-axes) and PKN-2 (different heatmaps correspond to 10x increase of RhoA-PKN-2 binding left to right as indicated). Colours correspond to the keys as shown in D, E.

Informed by the importance of Src perturbation in the ODE model, and observations concerning caveolae activation by Src in Boolean simulations, we next sought to establish the importance of the balance between Cav activation via sensing decreased membrane tension embedded in the positive feedback loop and bypassing direct protein-protein interaction activation. While for the Boolean formulation the only options were for the operation Src → Cav to be present or absent, with the ODE model we could tune the rate of this parameter simultaneously with the input of ‘polarised substrate stiffness’ to view the effect on retraction speed. Partitioning the parameter space domain into a 5×5 logarithmic square (Figure 4C) revealed the extent to which the coetaneous alteration of the initial polarised substrate stiffness value and the rate of active Src binding to inactive Cav1 effects rear retraction dynamics, and thus the extent to which the two disjunctive modes of activation can compensate for each other (Figure 4D, E). When the rate at which Src binds to (then activates) Cav1 was set at a low level, even extreme levels of stiffness polarisation were unable to potentiate a sufficient decrease in membrane tension to promote caveolae accumulation and subsequent RhoA activation (Figure 4C), preventing rear retraction. With negligible polarised substrate stiffness (i.e. on a near uniform substrate) potent binding and activation of Cav1 by Src caused rear retraction speeds of less than a third of the basal, co-operative levels (Figure 4C). When both values were set to low levels, retraction speed was negligible (red top left corner), while if both values are high, rapid recoil could occur (green bottom right corner, Figure 4D). More interestingly, a high degree of polarisation overcame relatively low Cav1 activation by Src but with a very slow transition time to steady state (Figure 4E), while high binding of Src to activate Cav1 was less able to compensate for low polarised substrate stiffness (Figure 4D). Taken together, these data indicate that the parameters we estimated for our initial model perturbations (polarised substrate stiffness – 100; Src-Cav1 binding rate -0.01) were well fitting, and that a low level activation of caveolae by Src in combination with a rigidity gradient is sufficient for efficient rear retraction (although other external factors may play a similar role, discussed later).

Next, we sought to elucidate the effect of the equilibrium of RhoA binding partners on rear retraction speed/efficiency. Three downstream binding partners were present in this model: DRF, ROCK1 and PKN-2. These interact with and act via several positive and negative effectors - LIMK, CPI-17, Cofilin, MLCP – to promote both F-actin alignment and MCL phosphorylation, which lead to acto-myosin contractility and rapid rear retraction. While the *in silico* knockdown of DRF, ROCK1 and PKN-2 all had some negative effect on retraction speed and efficiency in agreement with the suggestions of previous experimental findings, the theoretical over-expression of the proteins suggested that the balance of active RhoA binding was crucial for aligned actin and/or p-MLC: 10x increase in PKN-2 or DRF levels retarded retraction to negligible levels. Motivated by these findings, we sought to further ascertain the effects of different RhoA binding partners by simultaneously altering the rates of RhoA binding to DRF, ROCK1 and PKN-2 on a logarithmic scale, thus producing multiple 5×5 heatmaps (Figure 4F, G). This analysis revealed that potent RhoA-ROCK1 or RhoA-DRF binding alone were sufficient to lead to a steady state of high rear retraction, even with very low rates of RhoA binding to both its other two effectors. This compensatory effect of RhoA-ROCK1 binding was unsurprising considering ROCK1 phosphorylates MLC, activates CPI-17 and LIMK (which in turn inactivate the inhibitory MLCP and Cofilin respectively) and activates DRF which causes actin alignment, and thus contributed to all pathways leading to acto-myosin contractility. The compensatory effect of RhoA-DRF binding was more surprising; while the formin is the key actin organiser, it has no effect on p-MLC. It seems that even the lowest RhoA-ROCK1 binding rate is sufficient to promote enough p-MLC activity, while the F-actin alignment branch is more important to output dynamics of the model since it is an absolute requirement for both acto-myosin contractility and the rear retraction output itself (while p-MLC only effects the former). While this partitioning of parameter space revealed the combination of rates required for rapid rear retraction, the most interesting findings were with regards to the role of PKN-2. While any combination of RhoA-DRF and RhoA-ROCK1 binding with high enough rates were sufficient to promote cell rear retraction (top right and bottom left square of heatmaps in Figure 4F), RhoA-PKN-2 binding had a less positive effect on the steady-state speed of rear movement. Indeed, if the rate at which RhoA binds to PKN-2 was increased, retraction was in fact reduced for the same RhoA-DRF/RhoA-ROCK1 combinations (Figure 4F). While PKN-2 aligns F-actin and activates CPI-17, and thus acts to some extent in both the ‘Actin’ and ‘p-MLC’ pathway, this topology of the model suggested that it also had a role of sequestering active RhoA away from other important effectors (ROCK1 and DRF). Interestingly however, when RhoA-PKN-2 binding was too low (far left heat map, 4G) rear retraction takes an inordinately long time to initiate, while generally the higher the RhoA-PKN-2 binding rate, the quicker cells are able to organise their machinery to retract. Thus, while more ambiguous, our analysis has predicted that while DRF and ROCK1 had more obvious impact downstream of RhoA, PKN-2 binding is still important for quick responses to generate rear retraction, provided this was balanced and non-dominant. This further demonstrated the suggestion that, for cells expressing finite RhoA levels, the precise, regulated control of RhoA binding to partners is critical for efficient rear retraction.

### Further refinements of the ODE model: Ect-2 activation via CDK1

While the library of parameter scans of the initial model (Figure 3A) managed to corroborate key findings regarding the perturbation of variables which lead to efficient cell rear retraction, certain features of the simulated outputs were not entirely in-keeping with all previous experimental findings and thus the model could be further refined and adapted. In particular, while we have shown that caveolar sensing of localised membrane tension is important for localisation of Ect-2 to the rear domain and rapid rear retraction (Hetmanski *et al*., 2019), our *in silico* knockdown analysis here suggested an over-reliance of the whole system of caveolae, whereby at the 10% level of caveolae (Figure 3A), almost all retraction is abrogated. This result is more pronounced than *in vitro* findings, where Cav1 siRNA did indeed retard rear retraction experimentally, but to a lesser extent (Hetmanski *et al*., 2019). This effect is likely caused by caveolae acting as a ‘bottleneck’ in the formulation of the model, via which both Ect-2 was activated and Src (and subsequently p190RhoGAP) was inactivated. Based on an unbiased systematic GEF screen (Hetmanski *et al*., 2019), we included Ect-2 as the specific activator of RhoA in the rear retraction model(s), being activated directly downstream of caveolae. Phosphorylation of Ect-2 by CDK1 at T412 is required for GEF activity towards RhoA (Niiya *et al*., 2006), and so we postulated that this alternate activation of Ect-2 may make our model simulations more apt and physiologically plausible. We included the CDK activating protein (CAK) as an input and upstream activator of CDK1 based on literature evidence (Lolli and Johnson, 2005) (Figure 5A) while the rest of the model remained unchanged. The addition of these new variables/reactions had a small effect on unperturbed dynamics: rear retraction proceeded to a similarly high steady state value, albeit a few hundred seconds quicker (Figure 5B). With the inclusion of ‘external’ activation of Ect-2 alongside formation of caveolae by Src, we essentially now have two interactions serving a similar purpose within the model where the membrane tension dependent positive feedback loop is supported. The complete removal of either of these supporting interactions (e.g. the removal of CDK1 or Src) had a very minimal effect on rear retraction output dynamics, however the concomitant knockdown of both led to a complete loss of rear retraction (Figure 5C). Therefore if both activation of Ect-2 by CDK1 and caveolae by Src were at play during rear retraction events, individual experimental perturbation of either alone would have caused little difference for rear movement capacity. We performed experiments with CDK1 siRNA compared to control siRNA (knockdown efficiency 80%, Figure S5A) and found contrary to these model simulations that reducing CDK1 levels significantly retards global long-term cell movement (Figure 5C) and short-term rear retraction (Figure 5D) in 3D cell derived matrix (CDM), which was also supported by similar results using a specific CDK1 inhibitor (Figure S5B). These data suggest that CDK1 is important for rear retraction and may be a more relevant supporting interaction to directly activate Ect-2, and that the other interactions (caveolae activation by Src) should be amended to a lower rate accordingly. Upon this model alteration, CDK1 *in silico* knockdown now has a significant reductive effect on rear retraction in agreement with our experimental results in a switch-like manner (Figure 5E). Furthermore, with this iteration of the model, knockdown of Cav1 to 0-20% levels leads to a physiologically plausible but now not overestimated response on rear retraction, while Ect-2 remains a crucial protein for rear movement (Figure 5F). Perturbation of CDK1 to <20% in the model predicted a reduction of Ect-2 activity (via a loss of direct activation) and caveolae (more surprisingly, via the perturbation of the positive feedback loop). Indeed, CDK1 siRNA resulted in a loss of endogenous Ect-2 and Cavin-1 (a key component of caveolae) at the rear as predicted (Figure 5H). Furthermore, ratiometric FRET (Forster Resonance Energy Transfer) imaging showed that CDK1 knockdown led to a significant reduction in RhoA activity at the rear of migrating cells in CDM (Figure 5I), highlighting a specific, unexpected role for the usually cell cycle associated protein. These data suggest that activation of CDK1 is essential for efficient rear retraction and may have an important role in maintaining the membrane tension positive feedback loop to activate Ect-2 and subsequently RhoA.

Having established that CDK1 activity is important for efficient rear retraction, we next sought to compare the relative importance of Ect-2 activation by CDK1 and caveolae activation by Src directly. We used previously established specific inhibitors of CDK1 and Src as these permit a dose response to treatment with low and high levels of inhibition (as determined based on previous titration experiments (Horton *et al*., 2016; Jones *et al*., 2018)). In agreement with this current implementation of the model, CDK1 inhibition significantly reduced the capacity for cells to migrate in CDM at low or high doses (where at the high dose cells elongate and barely move), while Src inhibition alone only causes a significant retardation of cells at the high dose (Figure 5J, Movie S1). Furthermore, when both inhibitors were used at the same time at low doses, there was a small but significant reduction in migration speed compared to low CDK1 inhibition alone, in agreement with our earlier prediction regarding the targeting of both supporting interactions (Figures 5B, J, Movie S1). Altogether, these experimental data suggest we have improved on our initial kinetic model, supporting the new implementation with activation of Ect-2 by CDK1 and appropriately dampened (but still present) activation of caveolae by Src, and highlight the potential of CDK1 (potentially alongside Src) to abrogate cell rear retraction and global movement in 3D.

The model simulation also revealed a potential issue over the function of PKN-2. PKN-2 was important for the transition time to the steady-state of rapid rear retraction, but this variable had a negative effect on cell rear recoil dynamics if too dominant (conceptualised as sequestration of active RhoA away from other), such that *in silico* overexpression of PKN-2 or increased RhoA-PKN-2 binding rates reduces directional retraction to negligible levels. Experimental findings show that reduction of PKN-2 protein levels reduce cell speed and abrogate F-actin alignment at the rear (Hetmanski *et al*., 2019). The OR combinations of the reactions used in the model were therefore tested to see if the model could more faithfully recapitulate the experimental findings with multiple different parameter sets (Figure S6A, B). PKN-2 can clearly influence F-actin organisation in addition to contractility, but whether PKN-2 is a direct binder and organiser of F-actin is not clear (as shown by the dotted line in the schematics). We considered the possibility that PKN-2 has a pro-actin alignment role via synergistic activation. Therefore we made the following alterations in the model (Figure S6A): (i) PKN-

2 no longer binds to then subsequently aligns F-actin; (ii) DRF binding to F-actin relies on PKN-2 levels and (iii) PKN-2-CPI-17 binding is reduced tenfold (to balance PKN-2 activity). Incorporating these alterations, the unperturbed simulation reveals a slight increase in steady-state and little change in transition time thus showing the overall dynamics of the model are unchanged and remain in-keeping with experimental observations. More importantly, PKN-2 *in silico* knockdowns now had a big effect on the steady-state rear retraction speed and transition time to steady state (Figure S6B, left): levels less than 3% show negligible retraction, while levels 4-10% reveal unexpected dynamics where once retraction begins, it increased rapidly to an unstable local maximum before reducing slowly. Meanwhile with this new model formulation, *in silico* DRF knockdowns continued to display an almost linear, graded response and ROCK1 the characteristic switch-like response (as is seen in experimental knockdown/inhibition; (Paul *et al*., 2015; Hetmanski *et al*., 2019). This demonstrates that refining the model based on known activities of closely related proteins can enhance the plausibility, and thus the predictive power, of ODE based models.

### Stochastic individual cell rear simulations reveal more physiologically plausible predictions regarding critical substrate stiffness polarity

The deterministic nature of our modelling approach enabled us to capture average population dynamics, but not the behaviour of individual cells, which is highly heterogenous. Because the source of such heterogeneity is often difficult to understand, we sought to introduce stochasticity and randomness into our kinetic model to understand if the same given parameters could model the behaviour of individual moving cells within a population. *In vitro*, migrating cells exhibit a whole range of speeds and rear retraction efficiencies as well as protein expression and signalling levels. Starting from the initial model (Figure 3), we performed simulations using the stochastic τ-Leap method (of an irreversible implementation). Now, all the reactions relied on integer amounts of each variable, then provided there existed non-zero amounts of all variables involved in the reaction, a reaction would occur in the next time increment, randomly chosen based on the kinetic rate values and the concurrent variable levels (as is typical with mass action kinetics). Running 6 simulations to 20,000 seconds each, we saw that the rear recoil amount/speed in each simulation transitioned between ∼3000 and ∼6000 seconds to a high value of around 70%, before all simulations randomly fluctuated around this amount, ranging from a low of 55% to a high of 85%, showing some stochasticity but overall highly conserved activity between one another (Figure 6A, left). When comparing the average of these individual stochastic simulations with the deterministic simulation of the unperturbed ODE model (as in Figure 3), we saw that the outputs of the stochastic model were very similar to the steady state of the ODE model (Figure 6A, right). Overall however, the unperturbed simulations displayed reassuring agreement, while the stochastic model displayed a ‘subtle’ introduction of physiologically plausible rear retraction speed fluctuations.

**Figure 6:**
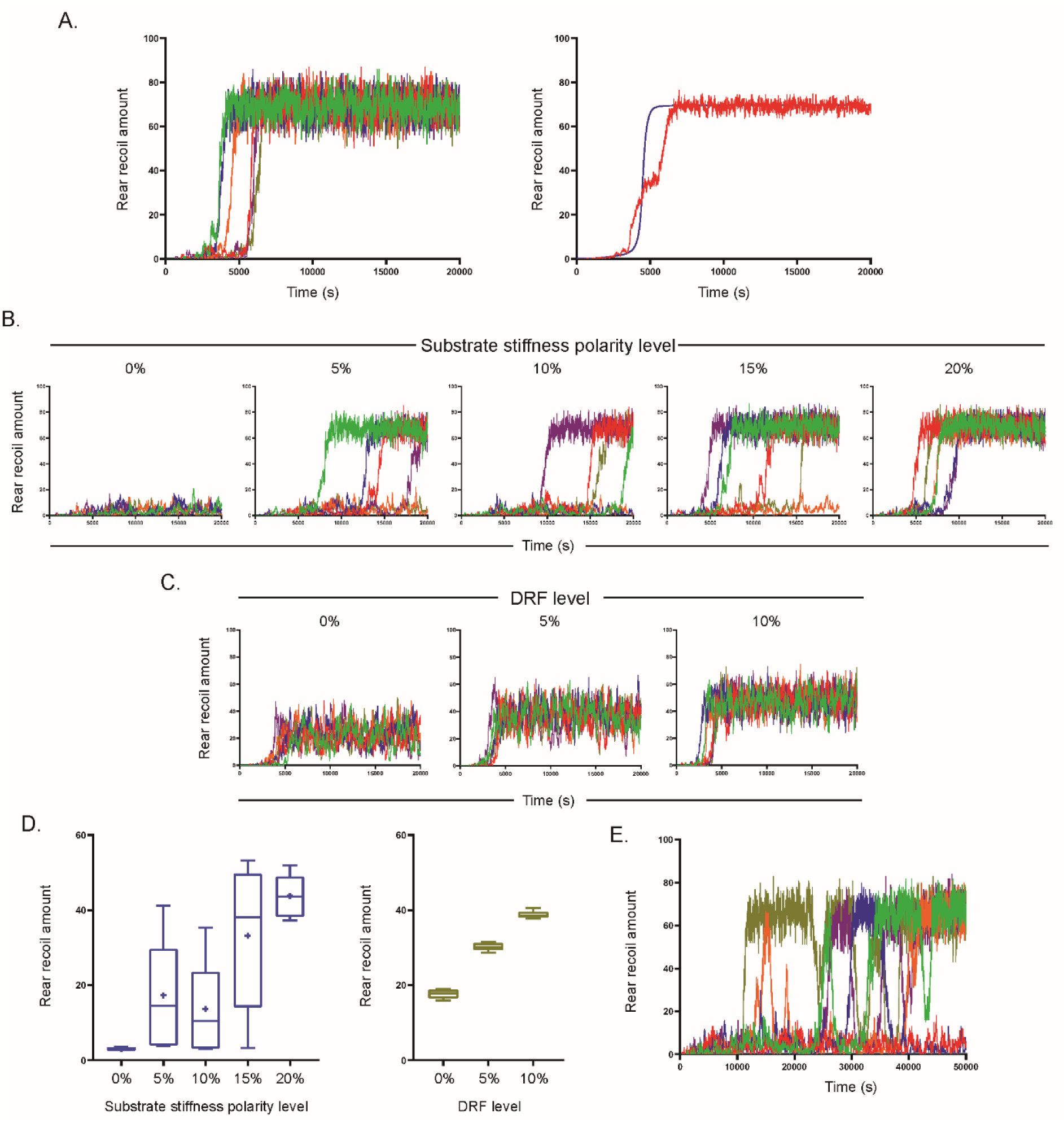
The stochastic rear retraction simulator makes more plausible predictions regarding individual cell behaviour. **A.** 6 stochastic simulations showing rear recoil amount with respect to time for the first 20,000s when run with exactly the same reactions and parameters as in the ODE model in Figure 3 (left) and the average of the 6 individual stochastic simulations (red line) in comparison with the previous deterministic curve (blue line) for rear retraction amount with the same parameters (right). **B, C.** 6 stochastic simulations showing the timecourse for the first 20,000s of rear retraction amounts with **B.** different substrate polarities, ranging from 0% (left) to 20% (right) of the unperturbed value and **C**. different DRF amounts, ranging from 0% (left) to 10% (right). **D.** Average rear retraction amount of all 6 cells across the whole 20,000s for each level of perturbation of left: substrate stiffness polarity; and right: DRF amount. **E.** 6 stochastic simulations showing the timecourse for the first 50,000s of rear retraction amount at the critical substrate stiffness polarity level (4%).

We next wanted to test perturbations of key model variables with the reactions/parameters as in Figures 3, 6A. In particular, *in silico* knockdowns in the ODE model showed that for certain variables there was a striking, switch-like response, while perturbation of other variables displayed more linearly/quadratically-dependent behaviour. We therefore performed similar perturbation analysis of our stochastic rear retraction simulator on two variables: the previous switch-like input of substrate stiffness polarity and the graded response to DRF amount. We performed 6 stochastic simulations each at values of stiffness polarity ranging incrementally from 0 to 20% for the first 20,000s (Figure 6A). With polarised stiffness removed (0%), all simulations displayed low retraction (<10%) throughout, while with high polarity (20%) simulations displayed similar dynamics to unperturbed simulations, where all 6 simulations reached the similar pseudo-steady-state by 10,000s. At intermediate levels of substrate stiffness polarity however (5, 10 and 15%), greater variability between the 6 simulated ‘cells’ was observed, whereby at all three of these polarity levels the transition time to the same high level rear recoil state was markedly erratic, while at least one cell was unable to reach the high state at all during the 20,000s at each polarity. The average rear recoil amount per simulation further highlighted this variability at intermediate stiffness polarity levels (Figure 6D, left), where average recoil at 5% polarity was in fact slightly higher than at 10%. Meanwhile the stochastic simulations at 0, 5, and 10% levels of DRF (Figure 6C) showed a near linear increase in average rear retraction speed dependent on DRF level (Figure 6C, D right), with numerical values and transition times highly analogous to the previous deterministic simulations of the ODE model (Figure 4B).

Motivated by the variability of different substrate stiffness polarity simulations (Figure 6D) alongside the identification of a critical level of stiffness polarity which cells were able to sense in the ODE model (Figure 4A) (above which – eventually – the rear retracted with similar speed to the unperturbed case, and below which retraction was negligible), we performed stochastic simulations at the integer value of substrate polarity nearest to the switch point (4%) and observed longer simulations (50,000s). At this critical substrate polarity level, all 6 simulations displayed pronounced variability both between simulations and for the same simulations at different times. Now the simulations all showed bistability, whereby rear retraction was either in a low state (<10%) or a high state (>60%) and transitioned rapidly between the two. Moreover, unlike with any of the ODE simulations (or the stochastic simulations when stopped after 20,000s), ‘cells’ did not remain in the high state of retraction, but instead bursts of high and low activity were observed, corresponding to physiologically plausible phases of rapid rear recoil interspersed with negligible directional movement. These data suggest that dynamics between the two models are similar when transition times to steady state are fast, but diverge when transition times are (perhaps physiologically implausibly) slow. This highlights when the stochastic rear retraction simulator will be of particular use as a predictive tool in future, and where it will most likely agree with deterministic findings.

### Stochastic simulations of the CDK1 model reveals outputs which agree with experimental observation of cells migrating in 3D extracellular matrix

Finally we sought to perform stochastic simulations with our refined ODE model where Ect-2 is activated by CDK1. Again, simulations of the unperturbed model showed close agreement with the previous ODE simulations during the first 20,000s with subtle stochasticity displayed (Figure S7). We therefore decided to test how stochastic simulations of perturbation of CDK1, Ect-2 and caveolae effect dynamics over a longer time course (50,000s). With deterministic simulations, varying CDK1 showed a switch like response (where at 20% rear recoil was negligible, but at 30% the high rear recoil steady state was reached). Reducing CDK1 levels to between 10% and 25% using the stochastic model showed similar switch-like responses for each individual simulation: all ‘cells’ either stayed in a low (<10%) rear retraction state throughout, or (at a seemingly random time) the retraction reached the high state (and remained in the pseudo-steady state). Incrementally increasing the CDK1 initial amount from 10% to 25% correspondingly increased the number of simulations which reached the high recoil state by 50,000s and decreased the transition time to this state, as shown by the average recoil levels at each CDK1 level (Figure 7E). Doubling initial Ect-2 amount from 10% to 20% showed an even more pronounced switch like response (Figure 7B), which was also much more robust and less variable (Figure 7E). Perturbation of caveolae in the CDK1 model meanwhile showed a more graded response and plausible fluctuation in rear recoil amount however much less variability in transition to pseudo-steady-state time (Figure 7C) and total average rear recoil amounts between simulations (Figure 7E), similar to previous deterministic simulations. Reassuringly, these simulations agree with previous experimental perturbation of all these key proteins via siRNA (Figure 5, (Hetmanski *et al*., 2019)), indicating that the stochastic simulation model may be the most accurate representation of 3D rear retraction.

**Figure 7:**
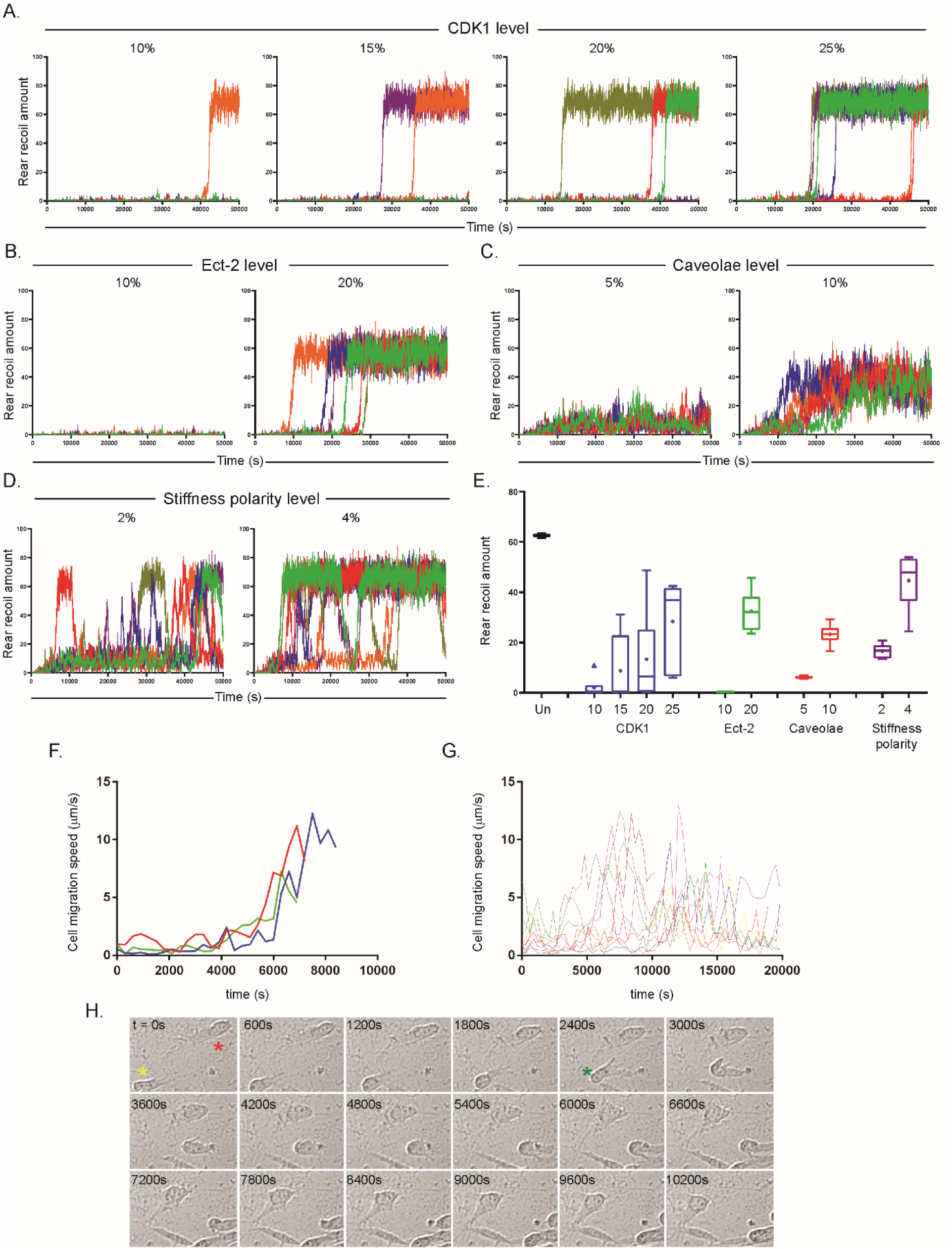
Stochastic simulations of the CDK1-included model recapitulates previous perturbation dynamics and predicts real cell behaviour in A-D. 6 stochastic simulations of the CDK1-included model showing the timecourse for the first 50,000s of rear retraction amounts with: **A.** different CDK1 amounts, ranging from 10% (left) to 25% (right) of the unperturbed value; **B.** Ect-2 amounts at 10% (left) and 20% (right), **B.** caveolae amounts at 5% (left) and 10% (right); and **D.** substrate stiffness polarity at 2% (left) and 4% (right). **E.** Average rear retraction amount of all 6 simulations across the whole 50,000s for unperturbed (denoted un), and each level of CDK1, Ect-2, caveolae and substrate stiffness polarity perturbation simulations. Note each individual simulation was averaged first and then the data of the 5 cells shown as a Tukey box plot, such that the spread corresponds to the spread between overall rear retraction amounts between different simulations (not the spread between values at different time points). **F.** Experimental automated tracking data of 3 A2780 cells starting from an unpolarised state, showing the instantaneous migration speed with respect to time, up until maximum rates of migration speed are reached. Cells were spread on CDM and imaged within 1 hour. **G**. Experimental automated tracking data of 10 randomly selected A2780 cells across 3 repeats showing instantaneous migration speed with respect to time across the whole time course or until cells go out of view. **H.** representative montage images of polarised A2780 cells every 600s across 10,800s, yellow star denotes a fast moving cell which undergoes a particular rapid phase of rear retraction at between t=2400s and t=3000s (denoted by green star), red star denotes a slower cell which has travelled much less distance in the same time period.

Finally, we sought to test our optimised stochastic CDK1 model in an environment which mimicked the environment of a cell derived matrix (CDM). We have seen previously that while CDMs are reasonably chaotic and show variability in substrate stiffness, cells often move in shallow stiffness gradients (Hetmanski *et al*., 2019). Translating this to our model, we tested simulations at low substrate stiffness polarity levels: 2% and 4%. At both these levels of polarity, simulations display bi-stability with bursts of high rear retraction interspersed with periods of lower directional movement at the back of cells (Figure 7D). Doubling the polarity of the substrate simply increases the amount of time each simulated ‘cell’ spends in the high recoil state without altering the bi-stable dynamics, while at both polarity levels reasonable ‘cell to cell’ variability is observed (Figure 7D, E). The stochastic cell simulator therefore predicts that, for unperturbed cells in a shallow gradient of rigidity (such as CDM), rear retraction is not a continuous, steady process but instead shows random bursts of fast retraction intertwined with periods of much less efficient rear contractility. Moreover it also shows that some cells can retract their rears significantly quicker than others. Interestingly, observing cell migration speeds with respect to time *in vitro* suggests both these findings may be biologically apparent (Figure 7F-H), thus underlining the value of developing a stochastic simulator in addition to the ODE approach.

## Discussion

By adopting different distinct modelling approaches to study the dynamics of directional rear retraction in 3-D non-uniform environments, we sought to satisfy two main aims: (i) to interrogate, further understand and inform future experiments regarding our biological system of interest, i.e. the signalling and mechanical hallmarks which lead to efficient cell rear contractility; and (ii) to analyse the value of Boolean, ODE and stochastic models on exactly the same system. Both of these lines of interrogation have been under-represented in the past: cell rear dynamics in 3-D remains a poorly elucidated area, while the adoption of multiple modelling approaches within the same study is not common.

Overall, the three-pronged modelling approach helped to further interrogate cell rear dynamics, whereby each model/simulation approach generated some interesting and novel ideas, while the increasing complexity involved in moving from Boolean logic to the stochastic simulation via the ODE model increased biological plausibility with each step. For example, the Boolean approach highlighted the near irrelevance of specific initial conditions, meaning ICs could be effectively ignored throughout the rest of the study using the more complicated kinetic formulations, and highlighted the quandary of Src/caveolae dynamics in controlling rear retraction. The ODE approach meanwhile made many population level predictions (Figures 3-5) and highlighted gaps and potential divergences away from prior experimental knowns such as regarding Ect-2 and PKN-2 dynamics. Finally the individual stochastic cell rear simulator revealed refined, more plausible predictions and increased agreement with previous *in vitro* data regarding the variability of individual migrating cells. In more general terms, the disparate modelling approaches seem to complement each other: Boolean logic interrogates general topology, deterministic ODE simulations are capable of a exploring a diverse range of reaction schemes which Boolean logic is unable to recapitulate, while the stochastic simulations finally offers the ability to predict the behaviour of individual ‘cells’ within a population. Moreover, with a ‘working’ ODE model, further subsequent analyses and perturbation analyses can be generated quickly and automatically. This suggests that, particularly in medium sized ∼20 variables systems where the ‘ideal’ modelling approach is unclear, using a multi-pronged approach as adopted here should be of value in other studies.

Between our three approaches, we have made many new predictions regarding the key events which lead to efficient rear retraction in fast moving cells and potential mechanisms to abrogate such potentially metastatic-inducing contractility, and we have highlighted a several potential future avenues of investigation. Specifically, the model points us to areas where we can increase our understanding of the system as a whole, for example elucidating the role of Src on caveolae/general rear dynamics using inhibitors/activators; testing the speed of directional rear retraction in different gradients of polarised substrate stiffness to ascertain further whether there exists a ‘critical gradient’ as predicted and implied by our experimental data; refining the roles of PKN-2 and DRFs in rear retraction.

Several predictions between the deterministic and the stochastic simulations are highly similar and conserved, for example with regards to the graded response of directional rear retraction speed in response to DRF knockdown (Figure 4B compared to Figure 6C). However, other predictions diverge, even when derived from the exact same reactions and rates, in particular when varying the input polarised substrate stiffness. Perturbation of our ODE model leads to the prediction that there exists a critical gradient of substrate stiffness which caveolae are able to ‘sense’. At values just above the ‘critical gradient’, the polarisation will be difficult to sense, hence directional retraction speed will remain at low levels for a significant amount of time, before eventually the gradient is interpreted and the rear begins to retract rapidly (in an ideal system). This is an unexpected but physiologically plausible prediction which may be tested using defined 2D gradients (e.g. poly-acrylamide substrates). Moreover, the deterministic simulations further predict that cells will move no faster in very steep gradients compared to moderately steep gradients of substrate stiffness, as in both cases a maximum velocity is reached. When translating our stochastic *in silico* predictions to the biological setting, exciting predictions emerge. At shallow/non-existent gradient levels of substrate stiffness polarity, little directional rear retraction will occur whereas, on substrates with a steep gradient, all cells will move with high, sustained rear retraction speeds. At intermediate levels however, cells will migrate with fast and slow phases of rear retraction speed, while overall some cells will move much more rapidly than others. This intermediate situation mirrors what we observe experimentally in a less defined 3D CDM, where the ECM is non-uniform and more chaotic (Figure 7G, H). This prediction clearly differs from the less plausible deterministic simulation forecast of a switch point where rears will either move (if above) or not (if below). The stochastic cell rear simulator has therefore significantly augmented our understanding of rear dynamics and the value of a rule-based model in addition to one built on ODEs is clear.

A particularly important abstraction of both our kinetic models is that all variables have a conserved total value bound between 0 and 100. This is vital for the predictions regarding certain variables sequestering other key variables away from more significant roles (e.g. PKN-2 with RhoA, CPI-17 with ROCK1). Furthermore, much of the variability of the outputs in the stochastic simulation approach are likely linked to this (bearing in mind all reactions depend on integer values of reactants). Considering the modelled domain is very tightly and specifically located to just the membrane bound region right at the rear of polarised migrating cells (such that when we talk of protein activity, we implicitly mean localised protein activity) it naturally follows that all proteins at play in our system will be bound to finite levels, which may be relatively low. In the absence of localised abundance data (which would be incredibly difficult to ascertain), we therefore decided to adopt this 0 – 100 levels approach. While this is certainly a viable approach and is akin to normalising protein/biophysical entity expression, it may be interesting to alter this 0-100 formulation in the future.

By refining the ODE model further (Figure 5) to include CAK/CDK1 activation of Ect-2, it was apparent that caveolae activation by Src may not be as vital as first suggested, but instead the essential *in silico* inclusion is an alternative means by which RhoA is activated at the rear (in addition to the decreased membrane tension feedback loop). Indeed, we saw that CDK1 inhibition had a stronger effect on cell migration than Src inhibition, and moreover that the combination of CDK1 and Src inhibition at low doses resulted in an additive slowing of cells compared to either inhibitor alone, as we predicted with our improved model. We also tested CDK1 perturbation on rear retraction, caveolae/Ect-2 localisation and RhoA activity and found this is vital for rear retraction via the membrane tension based positive feedback loop (despite not actually being implemented within said loop). Although CDK1 is more usually associated with control of cell division (Santamaría *et al*., 2007), interphase functions have been reported (Jones *et al*., 2018). Given the requirement for precise spatiotemporal control of Rho GTPases and the cytoskeleton in cell division and cell migration, it is interesting that some of the machinery is shared. While there has been much ongoing discussion surrounding the use of CDK1 inhibitors in cancer treatment to specifically perturb proliferation via cell cycle progression (Asghar *et al*., 2015), our results make the novel suggestion that CDK1 targeting may be able to decrease the metastatic potential of cells.

In conclusion, both deterministic and stochastic approaches were able to fit many of our known priors, making the possibilities of their future use in further interrogating cell rear retraction, and other spatial signalling contexts, extremely promising. Indeed, we previously successfully used the ROCK1/PKN-2 inhibitor to show the unexpected loss of localised caveolae accumulation as guided by the ODE simulations (Hetmanski *et al*., 2019). We hope that our ODE model and stochastic rear cell simulator will prove valuable tools in general cell migration and metastasis studies in the future.

## Materials and Methods

### Model Construction

The general starting point topology of all the models used, as shown in the wiring diagram in Figure 1A, was constructed based on our previous experimental findings (Hetmanski *et al*., 2019) and augmented by literature mining (Yang *et al*., 1998; Hamaguchi *et al*., 2000; Ohashi *et al*., 2000; Grande-García and del Pozo, 2008; Priya *et al*., 2015; Zimnicka *et al*., 2016). The starting point of the model remains unchanged, thus to avoid repetition, see (Hetmanski *et al*., 2019) model construction section for full details and references of all interactions included.

### Boolean formulation

In the signalling network (Figure 1A), each node corresponds to a variable, either a protein or a biophysical entity, and each directed edge corresponds to a different directed interaction between the nodes from and to which the edge is joining. Each node in the set of all nodes N is either in an active “ON” state or an inactive “OFF” state, denoted 1 and 0 respectively in a binary domain ∀ n ∊ N; t ∊ Z+ : n(t) = 0 or 1. Permissible reactions are activation or inhibition only, and are combined using the 3 Boolean operators AND, OR, and NOT, which are sufficient to represent any logical relationship. In this Boolean formulation, all reactions take the same amount of time to occur, t = 1, then given the set of initial conditions in which at least one node is initially active, ∃ n ∊ N : n(0) = 1, nodes became active (transition from 0 → 1), inactive (1 → 0), or remained the same (0→0 or 1→1) after each time increment depending on the state of upstream nodes and the interactions which the nodes are affected by. We took the input node ‘polarised substrate stiffness’ and the node ‘Ect-2’ as initially ON in our simulations, unless otherwise stated. These initial conditions (ICs) were chosen since the input node (substrate stiffness polarity) is a necessity to study our system of interest, while due to the positive feedback loops contained within the model, at least one other variable must be ON, therefore Ect-2 was chosen as the candidate due to the plausibility that the Ect-2 will generally be activated in the rear region without caveolae formation.

Simple reactions in the model, i.e. nodes with one directed input, were coded as single activations or inhibitions. Where protein nodes are affected by two upstream activators but no inhibitors (specifically DRF and CPI-17), interactions were taken as OR reactions due to the absence of any reported protein complex formation. Where biophysical entities are affected by two upstream activators (polarised contractility, decreased membrane tension, acto-myosin contractility and rear recoil), these interactions were implemented as AND reactions based on biological findings (e.g. that acto-myosin contractility requires aligned actin and phosphorylated myosin together). Other ambiguous relationships are dealt with explicitly in Figures 1 and 2.

### Boolean simulation and *in silico* knockout

All Boolean simulations were performed in CellNetAnalyzer (Klamt, Saez-Rodriguez and Gilles, 2007), a Matlab plugin. Simulations were either performed using synchronous updates for the first 40 time increments (where all possible reactions that can occur in a time increment will happen in the next time increment) or random asynchronous updates for the first 1000 time increments (where in every time increment at most one executable reaction will occur at random according to a uniform distribution) as explicitly stated. For synchronous updates, the automatically generated heatmaps for the activity of all variables in the model are presented. For asynchronous updates, 10 simulations were run and the heatmaps manually assessed for the production of summary bar graphs shown. *In silico* knockouts and subsequent node classification were performed by removing all edges which lead into a single node and observing the effects on the output variable rear retraction. *In silico* knockouts were performed individually for all nodes in the model as stated.

### Generation of random initial conditions

Random initial conditions (Figure 2) were created using the random number generator in Microsoft Excel (which generates a random number between 0 and 1), multiplied by 2 before having the floor function applied to allow only 0 or 1 as desired by Boolean logic. A 0/1 value was generated for all but the input node (kept at 1) and the output node (kept at 0), and each set was incorporated into synchronous simulations. This process was repeated 5 times.

### Kinetic model formulation and parameter values

The formulation and hallmarks that the kinetic model was built on, and the estimation of the parameter values used, was previously outlined in detail in (Hetmanski *et al*., 2019). Differences with respect to the earlier Boolean implementation of the model are that the decrease in membrane tension is a reversible AND type reaction which depends on both high polarised contractility and high recoil (as opposed to two irreversible reactions before); and polarised contractility is a reversible AND type reaction which additionally relies on high acto-myosin activity as well as polarised substrate stiffness (the input). All other reactions and parameter values remain the same; for justification see (Hetmanski *et al*., 2019). The values of the 10 most important parameters (as revealed by the parameter sensitivity analysis) were halved and doubled (Figure S4) to confirm the validity of our parameter value estimates, as shown by the general hallmarks of the simulations remaining similar despite strong perturbation.

### Initial conditions

For unperturbed simulations, the initial conditions (ICs) used were the input ‘polarised substrate stiffness’ set to the maximum 100, active Ect-2 set to 50, active Src set to 50, active Cofilin and MLCP set to 100, while the active form of all other is set to 0. The polarised substrate stiffness and Ect-2 ICs were taken for the same reasons as in the Boolean formulation section above (Ect-2 set to 50 such that some but not all Ect-2 is initially active, while the Ect-2 activator – Caveolae – can further activate Ect-2); Cofilin and MLCP were set to 100 since these are inhibitory variables with no explicit activators, such that the only way these can be active is when they start active, while Src was set at 50% active initially since the role of Src in the model is ambiguous as previously discovered in the Boolean section.

### Deterministic simulations

The model was built and simulated using Copasi software version 4.27 (Hoops *et al*., 2006). Deterministic simulations were performed using the LSODA method for the first 20,000 – 100,000s as indicated, solved with an interval size of 100s. For perturbations such as knockdowns, the initial amount of inactive form of each variable was edited, e.g. for the 10% knockdown, the inactive state was initially set at 10, such that the conserved moiety of the variable in question remains at 10 instead of 100 throughout simulations, while for the 10x overexpression simulations, the initial inactive levels were set to 1000. For the input polarised substrate stiffness and the inhibitors Cofilin and MLCP, the initial active amount was instead perturbed, while for Src and Ect-2 which were originally set at 50% active and 50% inactive, the perturbations of knockdown to 1%, 10% and 10x overexpression were performed by setting ICs to 0.5% active + 0.5% inactive, 5% active + 5% inactive and 500% active, 500% inactive respectively. These perturbations, as well as the coetaneous perturbation of multiple parameters in the model for Figure 4, were performed using the parameter scan function in Copasi. Throughout the main figures, only the rear retraction output is shown for clarity, however the active values of all variables in the model can be seen for the unperturbed simulation (Figure S3) to ensure all parts of the model work plausibly.

### Parameter sensitivity analysis

Parameter sensitivity analysis was performed using the built-in Copasi tool with Retraction/Recoil as the target and all rates in the model as the analysed objects. Scaled parameter analysis was run, such that the concurrent rates used in the model were factored in according to the apparent importance of each parameter, while the total of the scaled parameter sensitivity summed to zero.

### Generation of heatmaps

Heatmaps in Figure 4 were created using the parameter scan function, with a logarithmic alteration of the parameters of interest to cover 5 orders of magnitude. Steady states were manually determined as the values of rear retraction which are reached and remained at until the end of the simulations (either at 50,000 or 100,000s). Transition times to steady-state were taken as the time to the floor function of the steady state (e.g. for the real steady state of 68.745, the first time at which rear retraction > 68 is taken), and were to the nearest 100s since this was the interval time used in simulations. Heatmaps were created in Microsoft Excel, using a scaled 3-colour look up table (LUT).

### Stochastic simulations

Stochastic simulations were performed in Copasi version 4.27. All reactions in the previous kinetic models were first converted to irreversible. The stochastic τ-Leap method was used for simulation with Epsilon = 0.001, max internal steps set at 10000 and no random seed selected. Simulations were run to t = 20,000s with 20,000 intervals and an interval size of 1s, or to t = 50,000s with 25,000 intervals and an interval size of 2s. 6 simulations were manually performed (to take into account simulation variability while still being visible on one graph) per condition analysed, where computational time per simulation was ∼10s. Throughout, data was exported and plotted using GraphPad Prism. As with deterministic simulations, throughout the main figures, only the rear retraction output of the model is displayed.

### CDK1 siRNA long-term high-end widefield migration

A2780 human ovarian cancer cells were maintained in RPMI-1640 medium (Sigma-Aldrich) supplemented with 10% (v/v) foetal calf serum, 1% (v/v) L-Glutamine and (v/v) 1% Antibiotic-antimycotic (both Sigma-Aldrich), incubated at 37 °C in a humidified 5% (v/v) CO2 atmosphere as previously described. Knockdown of CDK1 was achieved with an individual siRNA (Dharmacon J-003224-15) using the nucleofection method and imaged 48h later. SiRNA efficiency was determined by standard SDS-PAGE western blot of lysates as described previously (Hetmanski *et al*., 2019), using the antibodies Mouse Cdc2 (CDK1) and Rabbit ERK1/2 (both Cell Signalling Technologies). Specific low/high inhibition of CDK1 was achieved using the inhibitor RO3306 (Sigma cat# SML056) 30 minutes prior to imaging at 1/10 μM; specific low/high Src inhibition was achieved using the inhibitor Saracatinib (AZD0530) 30 minutes prior to imaging at 0.3/3 µM; with DMSO used as a vehicle control. All cells were seeded sparsely in cell derived matrix (CDM, (Cukierman *et al*., 2001)) and imaged for 16h using on an Eclipse Ti inverted microscope (Nikon) with a 20x/ 0.45 SPlan Fluar objective and the Nikon filter sets for Brightfield and a pE-300 LED (CoolLED) fluorescent light source with imaging software NIS Elements AR.46.00.0. Point visiting was used to permit multiple positions to be imaged within the same time-course and cells were maintained at 37°C and 5% CO_2_. Images were acquired using a Retiga R6 (Q-Imaging) camera. 4-6 randomly chosen positions per cell were captured every 10 minutes over 16h. 5 randomly chosen cells per position (meaning 20-30 cells tracked per condition per experiment) were individually manually tracked using the ImageJ plugin MTrackJ every 3 frames (i.e. using 30 minute timepoint intervals).

### Live fluorescent microscopy

For rear movement, control/CDK1 siRNA cells were transfected by nucleofection with Lifeact-Emerald 24h before imaging (24h post siRNA nucleofection) and spread on CDM for 4 hours prior to imaging. Cells were imaged on a CSU-X1 spinning disc confocal (Yokagowa) on a Zeiss Axio-Observer Z1 microscope with a 63x/1.40 Plan-Apochromat objective. An Evolve EMCCD camera (Photometrics) and motorised XYZ stage (ASI) was used with 488 nm laser excitation. Images were captured using SlideBook 6.0 software (3i). Randomly chosen representative control/CDK1 siRNA cells were chosen and imaged for 5 minutes using point visiting while being maintained at 37°C. The distances moved by the rear of each cell during the 5 minute imaging interval were manually analysed using the line tool in ImageJ.

For ratiometric FRET RhoA activity, control/CDK1 siRNA cells were transfected by nucleofection with the Raichu-RhoA biosensor (Yoshizaki *et al*., 2003) 24h prior to imaging and imaged by spinning disc microscopy as above in the CFP-CFP, YFP-YFP and CFP-YFP channels. Average rear RhoA activity was determined using our previously established automated tool (Hetmanski *et al*., 2016), where the control/CDK1 siRNA conditions were normalised to the average FRET ratio of CDK1 knockdown cells to ameliorate experimental repeat variability.

### Ect-2/Cavin-1 staining and quantification

A2780 cells were cultured, spread on CDMs and fixed in 4% PFA as previously described (Hetmanski *et al*., 2019). Cells were stained with a Cavin-1 antibody (Abcam) and a Ect-2 antibody (Santa Cruz) and incubated with appropriate secondary. Cells were imaged using an SP8 G-STED microscope with appropriate settings, > 10 randomly chosen cell imaged across 2 experiments. For quantification, a 3-D surface plot with smoothing 3.0 was applied to each image (ImageJ), and the rear and rest of cell peaks manually identified under these conditions. For the box plot, the average of the rear peak and the rest of cell peak was calculated per cell, then the values (rear peak) / (average) and (rest of cell peak) / (average) calculated and plotted to give normalised rear and rest of cell staining values which were independent of raw intensity of staining.

### Automated cell tracking

A2780 cells were seeded on CDM and imaged within 1 hour of seeding, such that most cells were in situ but not spread or fully polarised yet (to try to get an approximate for time zero corresponding to the time zero used in the model). Cells were imaged on a widefield Nikon microscope, acquiring confocal images every 300s (5 minutes) for up to 20,000s. Cells were automatically tracked using Imaris software, where Wavelet software identified individual cells as different objects. Tracks were manually checked for accuracy. Instantaneous cell speeds were then automatically determined based on this tracking data, and 10 cells randomly plotted from 3 experimental repeats.

## Acknowledgments

We thank the Bioimaging Facility for their help with microscopy; microscopes used in this study were purchased with grants from BBSRC, the Wellcome Trust, and the University of Manchester Strategic Fund. We thank Egor Zindy for creating the automatic cell tracker tool. We thank Martin Humphries for providing the Src/CDK1 reagents. We thank Yitong Zhang for her assistance throughout the project.

## Supplementary Figures

**Figure S1:**
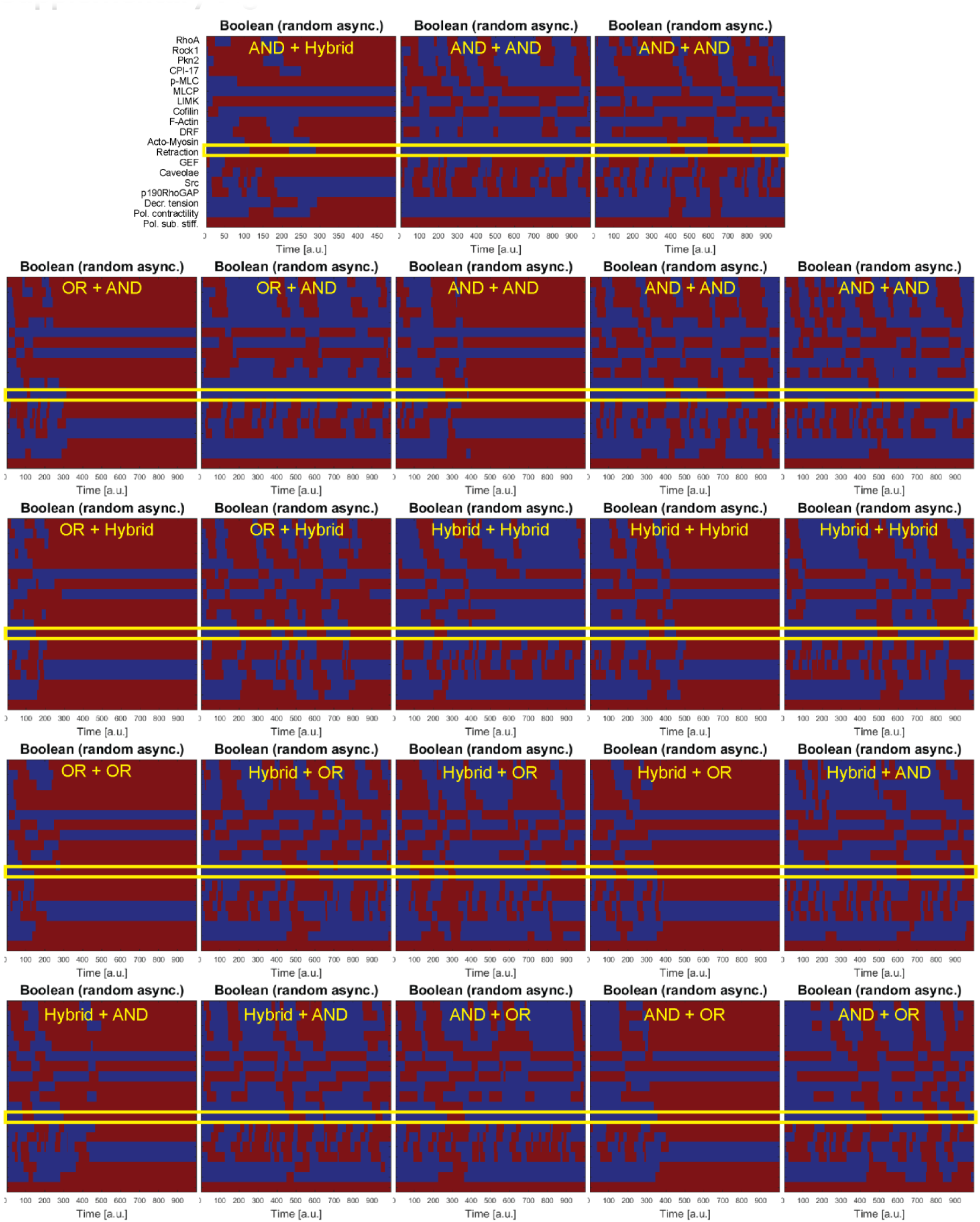
Random asynchronous Boolean simulations individual outputs. Between the 9 combinations of reaction schemes as in Figure 1B, and the three functional sets of behaviours of rear retraction: steady-state ON, cyclic bursts, steady state OFF, there is a total of 23 corresponding different outputs (i.e. OR + OR scheme only shows steady state rear retraction ON for all 10 stochastic simulations thus only one behaviour is possible whereas the AND + Hybrid scheme exhibits all three permissible behaviours within the 10 simulations). Heatmaps of one of each of these 23 behaviour/reaction scheme combinations are shown with rear retraction highlighted in the yellow box. Outputs were chosen randomly where appropriate.

**Figure S2:**
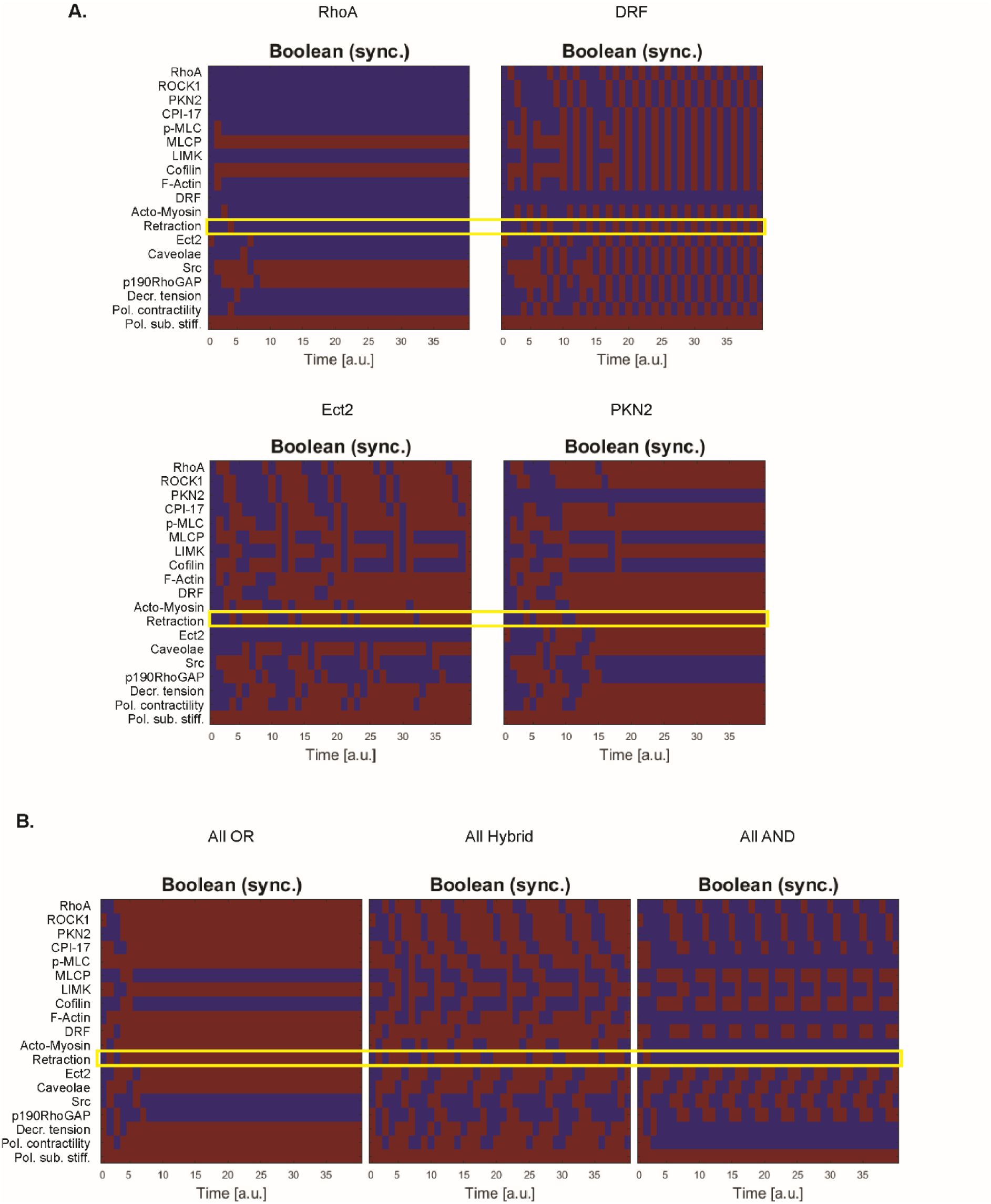
Activity heatmaps corresponding to *in silico* node knockouts and different reaction schemes for the same ICs. **(A)** Upon node removal (or constitutive activity for inhibitory variables), effects could be grouped into 4 functional sets as summarised in Figure 2D: rear retraction switches OFF, rear retraction switches to persistent oscillatory activity, rear retraction shows cyclic bursts of activity before reaching an ON steady state, and rear retraction remains with an ON steady state. Heatmaps of knockout of RhoA, DRF, GEF and PKN-2 respectively (all in the model without caveolae activation by Src, left in 2D) are shown to exhibit one of each of these behaviours with rear retraction output highlighted in the yellow box. **(B)** Heatmaps of the activity of all variables in the model with the reaction schemes OR + OR + OR (left), Hybrid + Hybrid + Hybrid (centre) and AND + AND + AND (right) given the same set of random initial conditions. Note the set of initial conditions used can be seen by the activity of all the variables at the initial time point (red – ON, blue – OFF).

**Figure S3:**
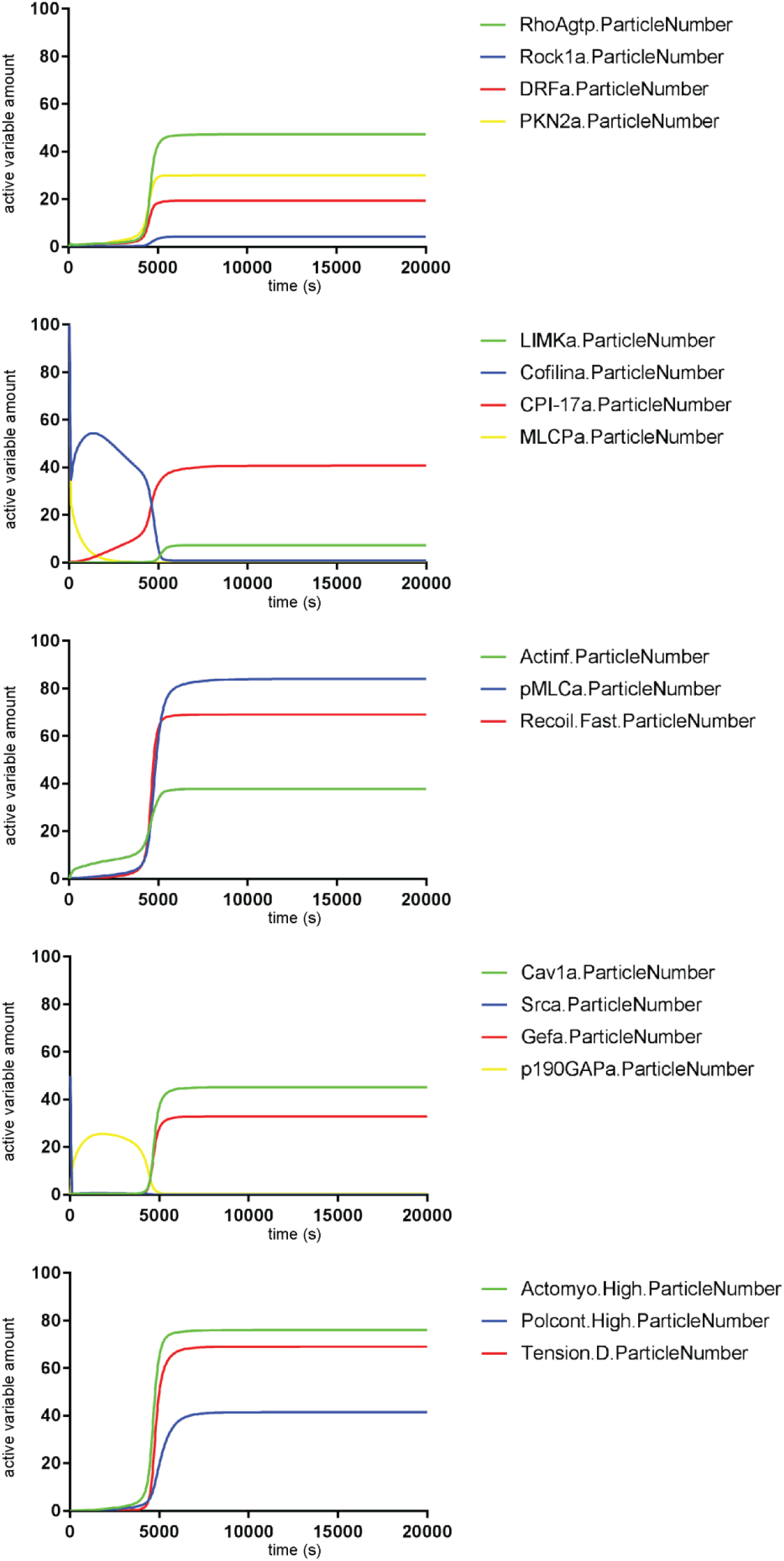
Timecourse curves for the activity levels of the active (unbound) form of all variables in the ODE model. Plots show the activity of the active form of all variables in the model during the first 20,000s. For visual convenience, curves are separated into 5 separate graphs: ranging from RhoA and effectors (top), to biophysical entities (bottom), colours or each curve corresponds to a variable as indicated. Note simulations are all corresponding to the unperturbed simulations in Figure 3A.

**Figure S4:**
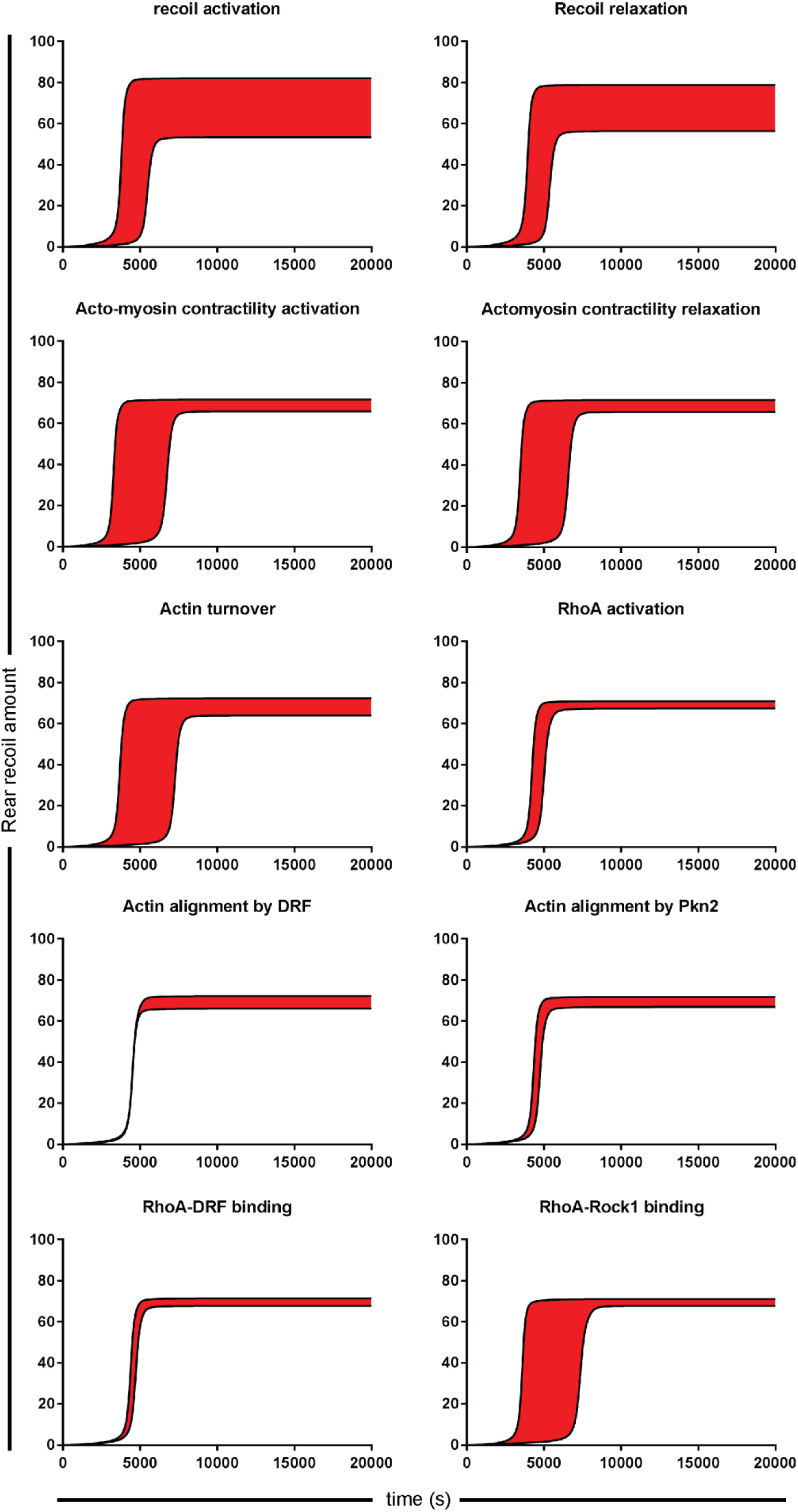
Range of rear retraction temporal dynamics given halving and doubling of the 10 most influential rates in the model. Time course curves of rear recoil amount given halving and doubling of the rate parameters (as shown above each graph) were plotted for the first 20,000 timepoints, then the region in between shaded red on the assumption that this whole region will be covered upon continuous alteration of the rate parameter between these two extremes.

**Figure S5:**
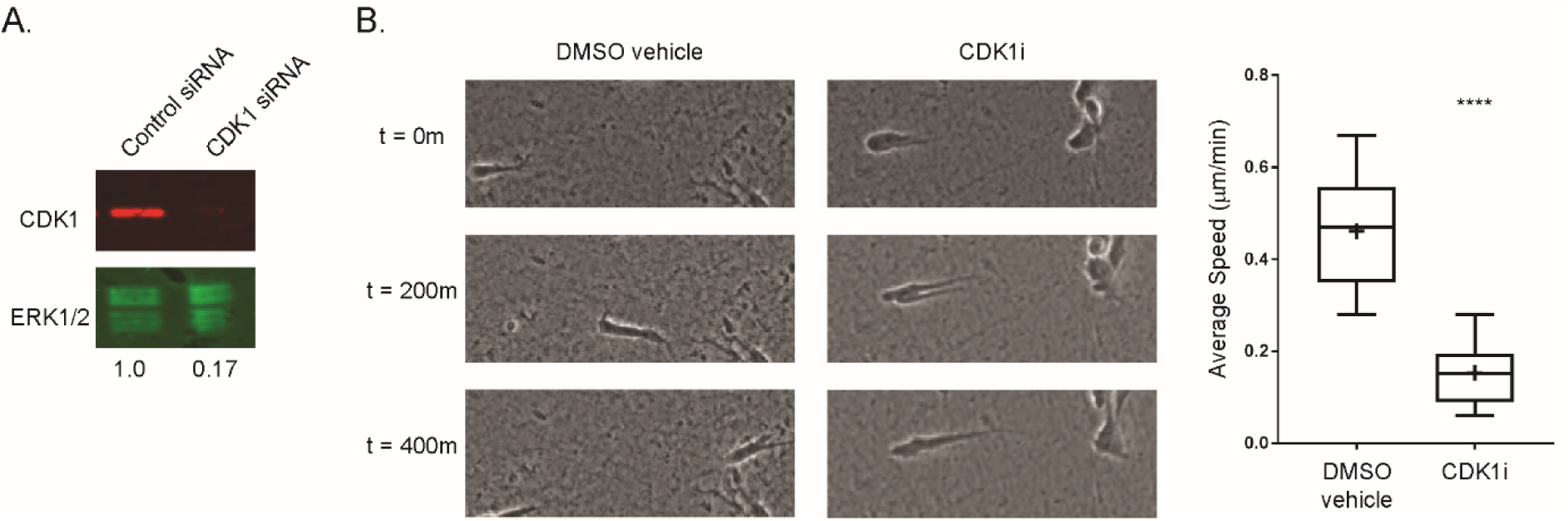
CDK1 siRNA efficiency and inhibition effect on cell migration. **A.** Efficiency of knockdown of CDK1 using individual siRNA (band intensity normalised to ERK1/2 loading shown below blots). **B.** Left: A2780 cells were treated with DMSO (left) or CDK1 inhibitor (right) and seeded in CDM and imaged by high-end widefield microscopy across 16 hours, representative individual cells shown across t = 400 minutes, Right: Quantification of average speed of control or CDK1 inhibited cells during 16h timecourse, (N = 30 cells across 2 repeats analysed per condition).

**Figure S6:**
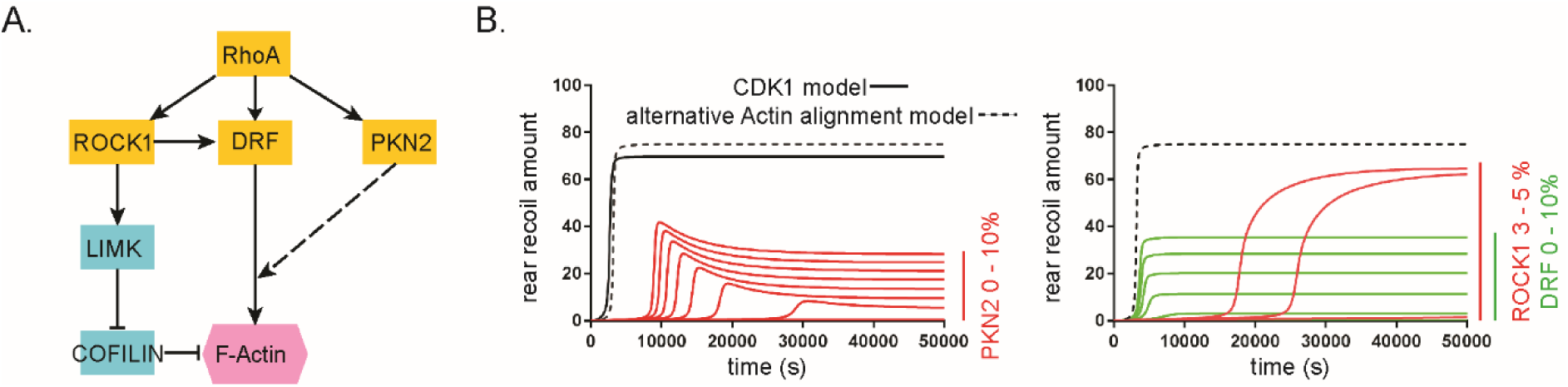
Alternative formulation of Actin alignment by PKN-2. **A.** Alternative formulation of model concerning RhoA signalling to aligned actin where F-actin alignment now requires coordinated activity of both DRF and PKN-2 together. **B.** Timecourse plots of rear retraction amounts in response to *in silico* knockdown/reductions in levels of PKN-2 between 0 and 10% (left), ROCK1 between 3 and 5% (red lines, right) or DRF between 0 and 10% (green lines, right).

**Figure S7:**
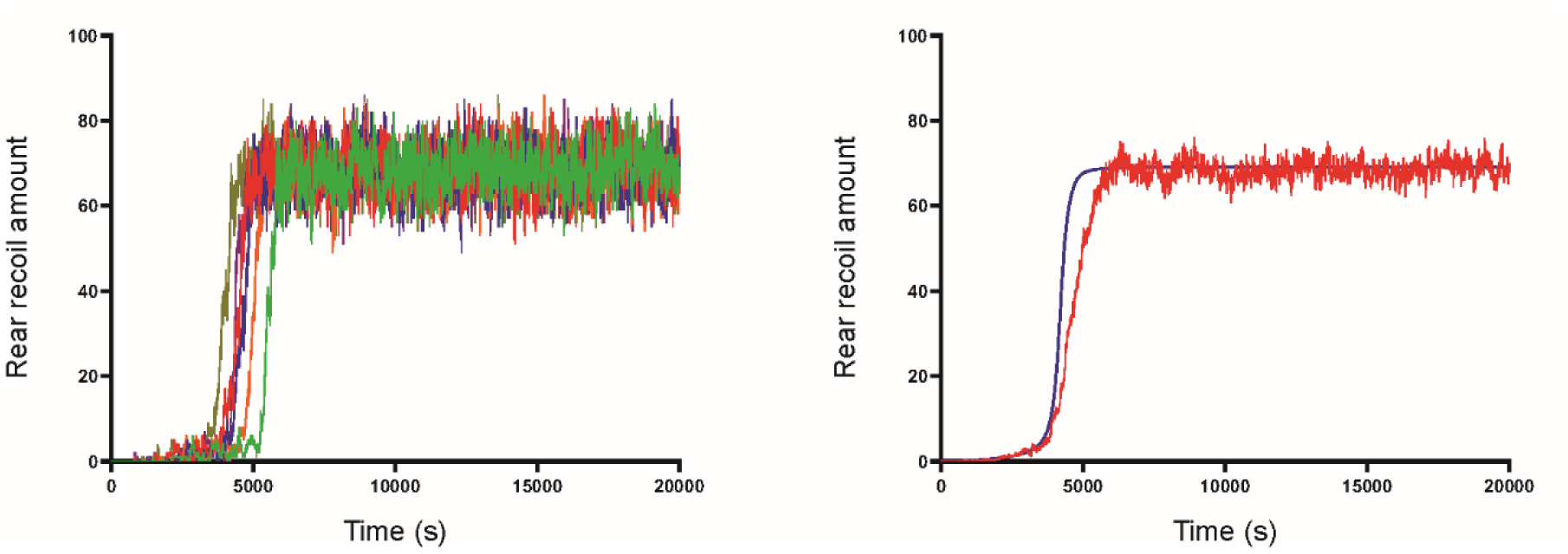
Stochastic simulation of the CDK1-included model. 6 stochastic simulations showing rear recoil amount with respect to time for the first 20,000s when run with exactly the same reactions and parameters as in the CDK1-included ODE model in Figure 5 (left) and the average of the 6 individual stochastic simulations (red line) in comparison with the previous deterministic curve (blue line) for rear retraction amount with the same parameters (right).

## Supplementary Movie

**Movie S1: Cell migration response to CDK1/Src inhibitor treatment.** Representative A2780 cells (moving left to right) migrating in 3D CDM over 7h treated with DMSO (control vehicle, top left); CDK1 inhibitor at 1 µM (top centre) or 10 µM (top right); Src inhibitor at 0.3 µM (bottom left) or 3 µM (bottom centre); or both CDK1 inhibitor at 1 µM and Src inhibitor at 0.3 µM together (bottom right).

